# HOX paralogs selectively convert binding of ubiquitous transcription factors into tissue-specific patterns of enhancer activation

**DOI:** 10.1101/871640

**Authors:** Laure Bridoux, Peyman Zarrineh, Joshua Mallen, Mike Phuycharoen, Victor Latorre, Frank Ladam, Marta Losa, Charles Sagerstrom, Kimberley A. Mace, Magnus Rattray, Nicoletta Bobola

**Author notes:** These authors contributed equally (Laure Bridoux, Peyman Zarrineh).

## Abstract

Gene expression programs determine cell fate in embryonic development and their dysregulation results in disease. Transcription factors (TFs) control gene expression by binding to enhancers, but how TFs select and activate their target enhancers is still unclear. HOX TFs share conserved homeodomains with highly similar sequence recognition properties, yet they impart the identity of different animal body parts. To understand how HOX TFs control their specific transcriptional programs *in vivo*, we compared HOXA2 and HOXA3 binding profiles in the mouse embryo. HOXA2 and HOXA3 directly cooperate with TALE TFs and selectively target different subsets of a broad TALE chromatin platform. Binding of HOX and tissue-specific TFs convert low affinity TALE binding into high confidence, tissue-specific binding events, which bear the mark of active enhancers. We propose that HOX paralogs, alone and in combination with tissue-specific TFs, generate tissue-specific transcriptional outputs by modulating the activity of TALE TFs at selected enhancers.

## Introduction

Gene expression programs instruct and maintain cell fate in embryonic development and adult tissue homeostasis. Transcription factors (TFs) control gene expression by binding to enhancers (Reiter et al., 2017; Spitz and Furlong, 2012). However, we still have no clear idea of how TFs select their precise sets of target enhancers. While TFs contain DNA binding domains which recognize DNA in a sequence-specific manner, these interactions are typically insufficient to direct a TF to its functional targets.

Transcriptional regulation is mediated by TFs working together, rather than in isolation. The widespread occurrence of collaborative TF binding is imposed by chromatin. A single TF cannot easily compete with nucleosomes to access DNA, but multiple TFs that recognize closely spaced binding sites can effectively displace nucleosomes and indirectly facilitate each other’s binding (Mirny, 2010; Moyle-Heyrman et al., 2011). Such indirect cooperativity can also result in TFs recognizing low affinity sites, i.e. sites that deviate from their optimal consensus *in vitro* (Farley et al., 2015). Recent observations indicate that TF cooperativity does not end at binding enhancers: clusters of enhancer-bound TFs concentrate co-activators and other nuclear factors via dynamic fuzzy interactions, driven by their intrinsically disordered regions (IDRs). IDRs function in molecular recognition and mediate the interaction with a diversity of regulatory proteins (Cumberworth et al., 2013; Staby et al., 2017) to promote the liquid-liquid phase transition associated with gene activation (Boija et al., 2018). Thus, the formation, on DNA segments, of regulatory complexes made of different combinations of factors, is key to activation of gene expression. These distinct combinations of TFs produce virtually inexhaustible flavours of gene expression and cell fate (Spitz and Furlong, 2012).

HOX TFs provide an ideal model to explain how TFs select their target enhancers to direct specific transcriptional programs *in vivo*. They contain a homeodomain (HD), a highly conserved DNA binding moiety shared by hundreds of TFs (Bobola and Merabet, 2017; Burglin and Affolter, 2016). HD display highly similar sequence recognition properties and bind the same core of four-base-pair sequence TAAT (Noyes et al., 2008), yet HOX TFs function to establish the identity of entirely different body parts along the antero–posterior axis of all bilaterian animals (Krumlauf, 1994; Pearson et al., 2005). In mammals, there are 39 *Hox* genes, classified into anterior (HOX1-2), central (HOX3–8), and posterior (HOX 9–13) paralog groups (Rezsohazy et al., 2015). HOX paralogs occupy sequential positions along the chromosome, which are faithfully maintained across evolution (Duboule, 2007). This translates into precise HOX expression codes at different levels of the antero-posterior axis, conferring specific spatial and temporal coordinates to each cell.

HOX association with three amino acid loop extension (TALE) HD TFs PBX, and PBX partner MEIS, is a widely accepted mechanism underlying HOX target specificity (Bobola and Merabet, 2017; Merabet and Mann, 2016; Selleri et al., 2019). HOX-TALE cooperativity increases the affinity and sequence selectivity of HOX TFs *in vitro* (Merabet and Mann, 2016). *In vivo*, HOXA2 extensively binds with TALE TFs (Amin et al., 2015) and Ubx and Hth (fly homologs to vertebrate central HOX and MEIS respectively) co-localize in active nuclear microenvironments, suggesting that their interaction may be critical to trigger phase separation (Tsai et al., 2017). Interestingly, Hox binding selectivity can be observed in the absence of TALE TFs, and is strongly associated with chromatin accessibility (Porcelli, 2019). Although the concept of HOX and TALE interaction is long established, we still understand relatively little about the extent and functional significance of HOX-TALE association *in vivo*, where compaction of DNA into chromatin and the distribution of sequence-specific TFs (cell-specific and tissue-specific, but also ubiquitous) can considerably affect TF binding to DNA. Also, how the association with fairly ubiquitous proteins eventually translates into HOX paralog-specific transcriptional outputs *in vivo*, remains unclear.

To understand how HOX TFs execute their specific functions to impart different segmental identity *in vivo*, we compared binding of HOXA2 and HOXA3, an anterior and a central HOX proteins, in the physiological tissues where these TFs are active. Branchial arches (BA) are blocks of embryonic tissues that merge to form the face and the neck in vertebrates. The second and third branchial arch (BA2 and BA3) are the main domains of HOXA2 and HOXA3 expression respectively, and the embryonic areas most affected by inactivation of *Hoxa2* and *Hoxa3* in mouse (Gendron-Maguire et al., 1993; Manley and Capecchi, 1995; Rijli et al., 1993). We find that HOXA2 and HOXA3 occupy a large set of high-confidence, non-overlapping genomic regions, that are also bound by TALE TFs. We identify three main determinants of HOX paralog-selective binding, resulting in high-confidence cooperative HOX-TALE binding at different genomic locations: recognition of unique variants of the HOX-PBX motif, differential affinity at shared HOX-PBX motifs and, additional contribution of tissue-specific TFs. We propose that HOX paralogs operate, alone and in concert with tissue-specific TFs, to switch on TALE function at selected enhancers.

## Results

### HOXA2 and HOXA3 control diverse processes by targeting different regions of the genome

HOX TFs direct highly specific gene expression programs *in vivo*, but recognize very similar DNA sequences *in vitro*. However, it remains to be determined if HOX specificity of action reflects specificity of binding across the genome *in vivo*, i.e. the binding of paralog HOX TFs to distinct target regions. To establish this, we compared HOXA2 and HOXA3 binding profiles in their physiological domains of expression in the mouse embryo. BAs display an antero-posterior gradient of HOX expression, which replicates *Hox* gene positions on the chromosome (Fig. 1AB): BA1 does not express any *Hox* gene, BA2 expresses *Hox2* paralogs, BA3 *Hox3* paralogs, etc. We previously characterized HOXA2 binding in BA2 (Amin et al., 2015); here, we profiled HOXA3 binding in BA3-4-6 (hereafter referred to as posterior branchial arches, PBA), the embryonic tissues immediately posterior to the BA2 (identified by the expression of *Hox* paralogs 3-5, Fig. 1AB). Using a HOXA3-specific antibody (Fig. 1-Supplemental Fig. 1A), we identified 848 peaks with fold enrichment (FE) ≥10, which largely contained a second biological replicate (Fig. 1-Supplemental Fig. 1B). TALE TFs (PBX and MEIS) display cooperative binding with HOX and increase HOX binding specificity *in vitro* (Merabet and Mann, 2016). De novo motif discovery (Heinz et al., 2010) identified HOX-PBX recognition sequence as the top enriched motif in HOXA3 peaks and uncovered MEIS binding site in the top three sequence motifs (Fig. 1-Supplemental Fig. 1C). HOXA3 recognition sites in PBA correspond to HOXA2 motifs in BA2; moreover, the distribution of HOX-PBX motifs is comparable across HOXA2 and HOXA3 peaks. HOX peaks without a canonical HOX-PBX consensus motif, contain potential low affinity variants of HOX-PBX sites (Fig. 1-Supplemental Fig. 1D-F). The occurrence of high affinity sites (perfect matches) positively correlates with peak FE, and is highest in top HOXA2 and HOXA3 peaks. Low affinity sites (1 mismatch) show the opposite trend and occur with higher frequency in lower confidence binding events (Fig. 1-Supplemental Fig. 1D-F).

**Figure 1.**
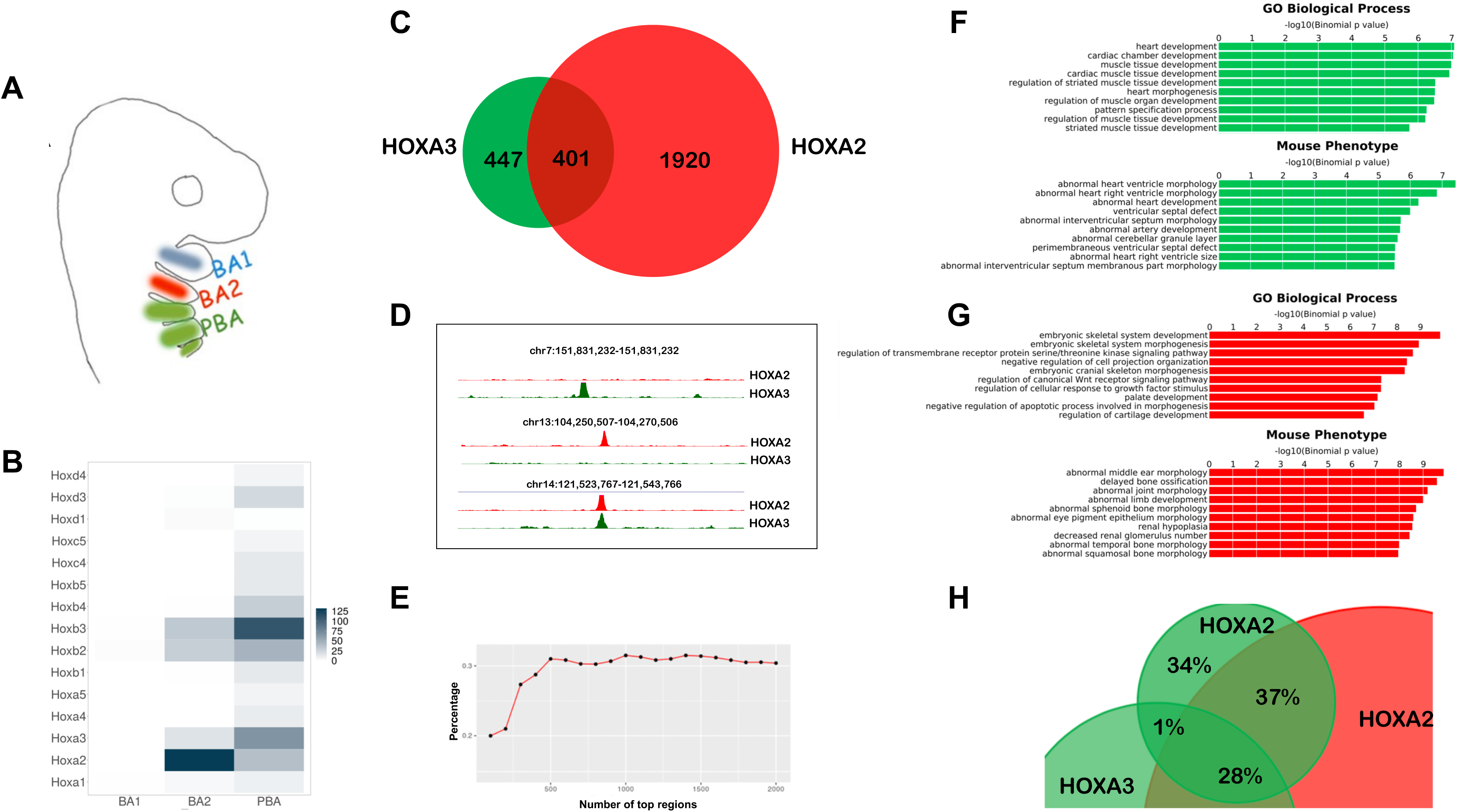
HOXA2 and HOXA3 control diverse processes by targeting different regions of the genome *in vivo*. A. BA organization in mammals. BA3-6 are collectively indicated as PBA. The same colour code (BA2 red, PBA green) is used throughout the manuscript. B. Heatmap of *Hox* expression in E10.5 mouse BA1, BA2 and PBA, based on the normalized expression values count per million (CPM)(Losa et al., 2017). C. Overlap of HOXA3 binding in PBA and HOXA2 binding in BA2 (200 nt summits, overlap at least 1 nt). Only peaks with FE>10 are considered. D. UCSC tracks (mm9) of HOXA3 (green) and HOXA2 (red) specific and shared peaks. E. Overlap (%) of increasing numbers of top HOXA2 and HOXA3 peaks (ranked by FE). High-confidence peaks show the smallest overlap. FG. GREAT analysis of HOXA3- (F) and HOXA2- (G) specific peaks (non-overlapping, green and red bars respectively) shows association with genes involved in different biological processes and whose mutations generate different phenotypes in mouse. The length of the bars corresponds to the binomial raw (uncorrected) P-values (x-axis values). H. HOXA2 binding in PBA. Overlap of HOXA2 summit regions in PBA (FE >10, green) with HOXA2 summit regions in the BA2 (red) and HOXA3 summit regions in the PBA (green); same rule as in C. HOXA2 binding locations are similar in BA2 and PBA.

We overlapped HOXA2 binding in BA2 with HOXA3 binding in PBA. About half of HOXA3 peaks are contained in the larger HOXA2 datasets (Fig. 1CD). When comparing the same number of peaks for both datasets, ranked by FE, we observed an increasing overlap at lower confidence peaks (Fig. 1E), suggesting that HOXA2 and HOXA3 select different sites when binding with higher affinity and are more promiscuous at lower binding levels. Functional association of HOXA3-specific peaks in PBA and HOXA2-specific peaks in BA2 (McLean et al., 2010)(Fig. 1FG) highlights distinct biological processes and mouse phenotypes, including abnormal middle ear, sphenoid, temporal and squamosal bone morphologies, whose morphogenesis is controlled by HOXA2 (Gendron-Maguire et al., 1993; Rijli et al., 1993). In contrast HOXA3-specific binding is almost exclusively associated with heart and cardiac muscle development and cardiovascular phenotypes, consistent with the role of HOXA3 in the formation of the main arteries (Manley and Capecchi, 1995, 1997) (Fig. 1F). These observations are in line with HOX functional specificity and indicate that in their physiological domains of expression, HOXA2 and HOXA3 bind in the vicinity of, and potentially control, genes involved in very different processes. *Hoxa2* expression displays a sharp anterior border between BA1 and BA2 and expands in the more posterior PBA (Fig. 1A; Fig. 4A). We profiled HOXA2 binding in PBA to understand if HOX-specific binding is determined by differences in the BA2 and PBA chromatin environment. We found that HOXA2 peaks in PBA very rarely overlap with HOXA3 ‘only’ peaks in the same tissue (1% overlap), but are largely contained in the pool of HOXA2-specific binding in BA2 and ‘common’ HOXA2 and HOXA3 binding events (Fig. 1H). This argues against differences in chromatin accessibility being a main determinant of HOX binding. In sum, analysis of HOXA2 and HOXA3 ChIP-seq in their respective domains of expression indicates that different HOX TFs control diverse and specific processes by targeting different regions of the genome *in vivo.* Tissue-specific chromatin accessibility does not appear to be a major determinant in HOX paralogs’ target site selection.

**Figure 2.**
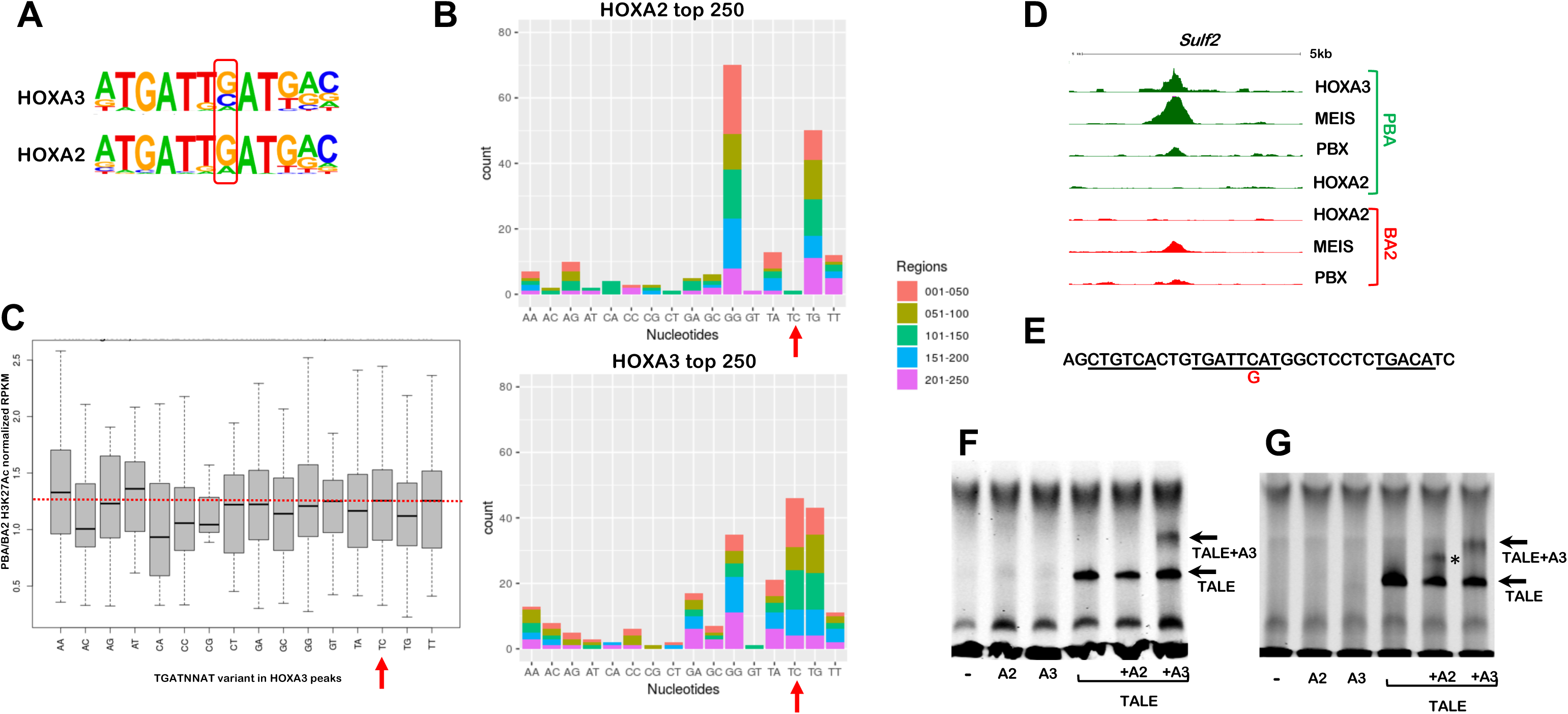
HOXA2 and HOXA3 select variants of the HOX/PBX motif. A. Homer detects different variants of the HOX-PBX motif in top 250 HOXA2 and HOXA3 peaks, with a G/C (HOXA3) or mainly a G (HOXA2) in the second variable position. B. Occurrence of HOX-PBX motif variants (all permutations of the variable nucleotides in TGATNNAT) in top 250 HOXA2 and HOXA3 peaks (ordered into 50 region bins). The TGATTCAT motif (red arrows) is among the most enriched variants in HOXA3 peaks but does not virtually occur in HOXA2 peaks. C. Box plot of global H3K27 acetylation levels (PBA/BA2 ratio) at HOXA3 peaks containing different TGATNNAT variants. HOXA3 peaks containing the TGATTCAT variant are associated with increased enhancer activity in PBA (red line). D. UCSC tracks with HOXA3, HOXA2, PBX and MEIS binding profiles in BA2 (red) and PBA (green) at the *Sulf2* locus, containing TGATTCAT. No HOXA2 binding is detected in BA2 or PBA. E. Sequence of HOXA3 peak summit in D, corresponding to the probe used in F. The TGATTCAT motif (underlined) is flanked by two MEIS motifs (also underlined); the C→G substitution tested in G is indicated in red. F. HOXA3 can selectively bind the *Sulf2* probe in complex with PBX/MEIS. Incubation of the *Sulf2* probe with TNT reticulocyte expressing HOXA2, HOXA3, MEIS/PBX, HOXA2/MEIS/PBX or HOXA3/MEIS/PBX. MEIS/PBX bind the *Sulf2* probe in combination (arrow). Addition of HOXA3 to the probe results in the formation of a complex only in the presence of PBX/MEIS (arrow). No complex is formed when PBX/MEIS are co-translated with HOXA2. G. Same experiment as in F, using a mutant *Sulf2* probe (the nucleotide substitution is shown in E). HOXA2 can bind the mutant probe in combination with MEIS/PBX (asterisk), similar to HOXA3 (arrow).

**Figure 3.**
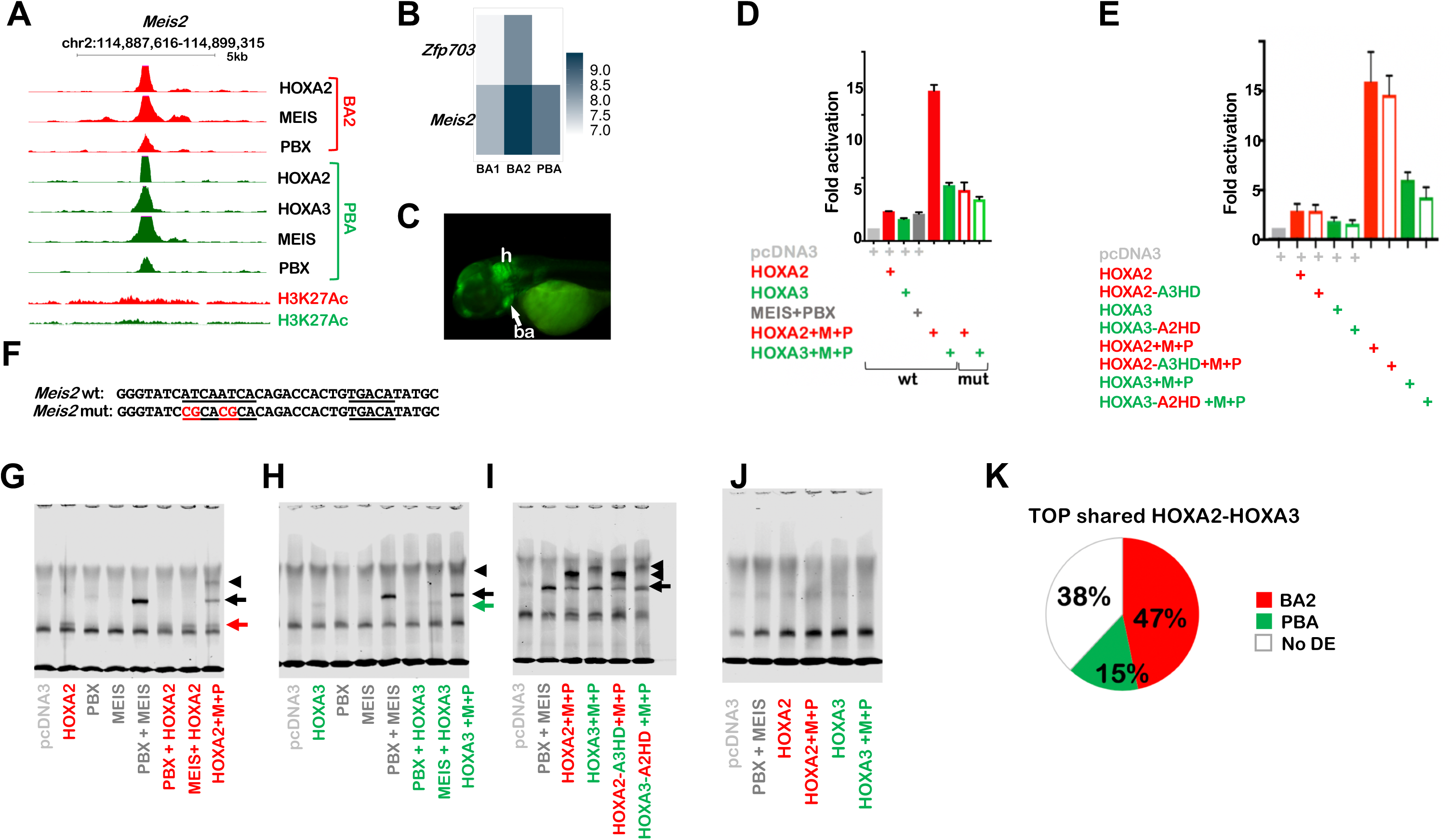
HOXA2 control of target enhancers. A. UCSC tracks of HOXA2, HOXA3, PBX, MEIS binding and H3K27 acetylation profiles in BA2 (red) and PBA (green) at the *Meis2* locus. Strong HOX and TALE binding is observed in both tissues, with higher acetylation levels in BA2. B. Heatmap shows *Meis2 and Zfp703* expression in E11.5 mouse BA1, BA2 and PBA, based on the normalized expression values CPM (Losa et al., 2017). C. *Meis2* enhancer is active in the hindbrain (h) and the BAs (ba, arrow) of developing zebrafish, which correspond to *Meis2* expression domains in mouse (Amin et al., 2015). The enhancer sequence spans the 200nt summit of HOXA2 peak in A. D. Luciferase activity driven by *Meis2* enhancer co-transfected with *Hoxa2* (red bar) or *Hoxa3* (green bar) in combination with *Meis2* and *Pbx1a* expression vectors in NIH3T3 cells. The combination of *Hoxa2* with *Meis2* and *Pbx1a* results in the highest activation. Changing the HOX-PBX site (empty bars, mutant sequence in F) reduces HOX-TALE activation. E. Luciferase activity driven by *Meis2* enhancer co-transfected with *Hoxa2-a3HD* (red empty bar) or *Hoxa3-a2HD* (green empty bar) and *Meis2* and *Pbx1a.* Values shown in DE represent fold activation over basal enhancer activity and are presented as the average of at least two independent experiments, each performed in triplicate. Error bars represent the standard error of the mean (SEM). F. Sequence of *Meis2* wild-type and mutant probe. HOX-PBX (reverse) and MEIS motifs are underlined. Nucleotide substitution in the HOX-PBX site are shown in red. G-J. Incubation of the *Meis2* probe with TNT reticulocyte expressing HOXA2, HOXA3, MEIS/PBX, HOXA2/MEIS/PBX or HOXA3/MEIS/PBX as indicated. G-H. HOXA2 (G, red arrow) and HOXA3 (H, green arrow) weakly bind the *Meis2* probe. MEIS and PBX bind DNA together (black arrow). Addition of HOXA2 results in a trimeric protein complex (arrowhead); the intensity of the MEIS/PBX complex is reduced (black arrow). Addition of HOXA3 results in a higher complex (arrowhead), but without affecting the intensity of the MEIS/PBX dimeric complex (black arrow). I. Swapping HOXA3-HD with HOXA2-HD does not improve the ability of HOXA3 to form a ternary complex with PBX and MEIS, and does not decrease HOXA2 binding with MEIS and PBX (arrowheads). Adding HOXA2 (or HOXA2-A3HD) results in higher intensity of the trimeric complex and lower intensity of TALE dimeric complex relative to HOXA3 (or HOXA3-A2HD), as observed in G-H. J. *Meis2* mutant probe (sequence in F) does not interact with HOX and/or TALE. K. Top HOXA2 and HOXA3 overlapping peaks (total of 60 intersecting top 250 HOXA2 and HOXA3 peaks) are more frequently associated with genes with higher expression in BA2 (red) relative to PBA (green). The white portion of the pie chart refers to genes that are not differentially expressed (no DE). Gene association is based on GREAT standard association rules; expression levels are extracted from E11.5 RNA-seq (Losa et al., 2017).

**Figure 4.**
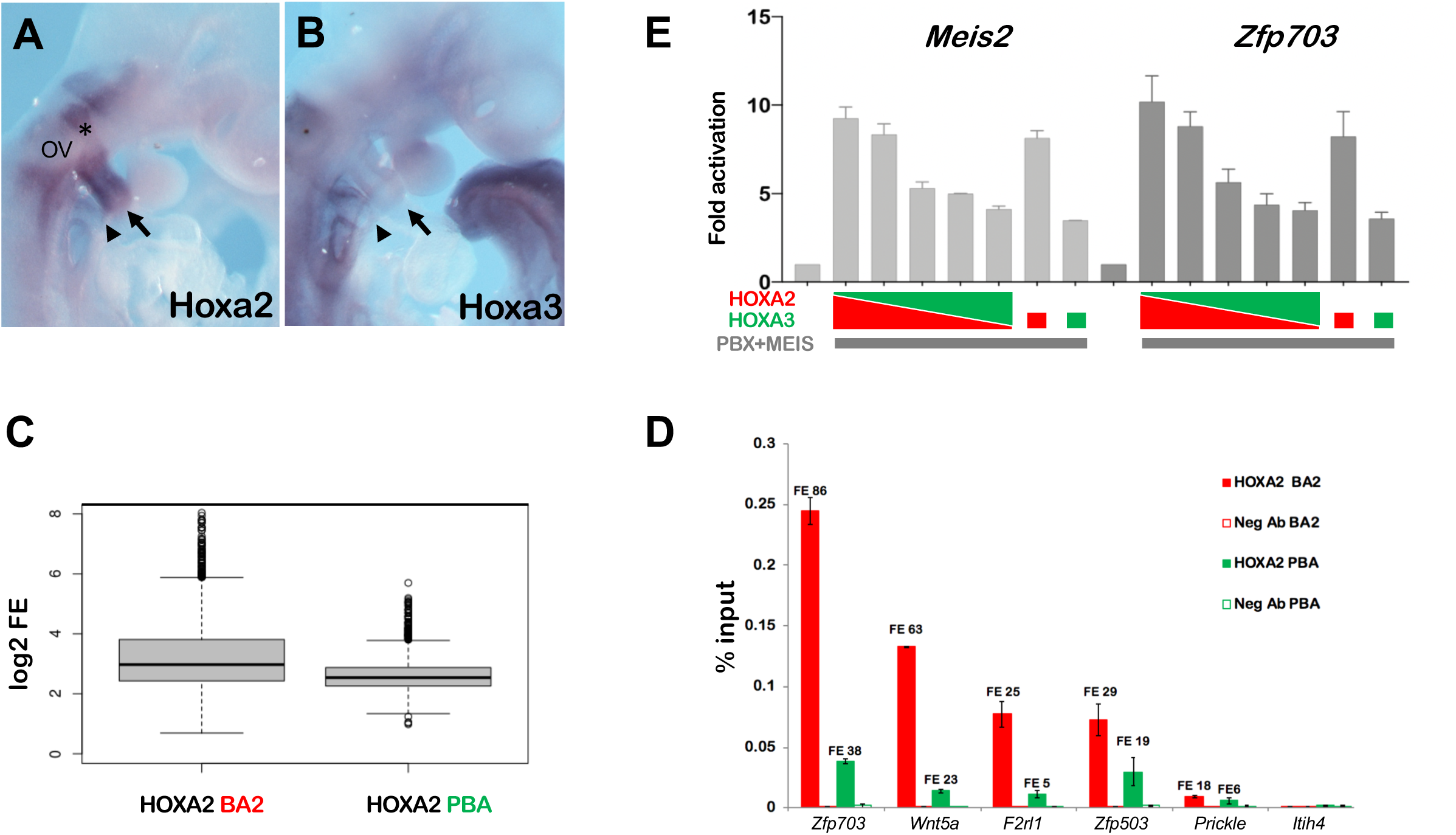
AB. *In situ* hybridization on E9.5 embryos, using *Hoxa2* (A) and *Hoxa3* (B) probes. A. *Hoxa2* is highly expressed in the neural crest migrating from rhombomere 4 (asterisk) to the BA2 (arrow). The portion of neural crest migrating just below the otic vesicle (OV) into the BA3 (arrowhead) is also *Hoxa2*-positive. B. *Hoxa3* is expressed in the BA3 (arrowhead). C. Boxplots of FE of HOXA2 peaks in BA2 and PBA. D. Comparison of HOXA2 binding in BA2 (red bars) and PBA (green bars) by ChIP-qPCR. Enrichment of each region following immunoprecipitation with HOXA2 and IgG negative control antibody (Neg Ab) is calculated as percentage input; numbers indicate the corresponding FE values in HOXA2 ChIP-seq (BA2 and PBA). Peaks are labelled by their closest genes. *Itih4* is a negative control (unbound region). Values represent the average of duplicate samples, and error bars indicate the SEM. D. Luciferase activity driven by *Meis2* and *Zfp703* enhancers co-transfected with expression vector for *Hoxa2* or *Hoxa3*, alone, or at diverse ratio of *Hoxa2* to *Hoxa3* (3:1; 2:2; 1:3) as indicated. All samples, except the negative control, contain *Hox* in in combination with *Meis2* and *Pbx1a* expression vectors. For both enhancers, luciferase activity decreases as *Hoxa2* is progressively replaced by *Hoxa3.* Values represent fold activation over basal enhancer activity and are presented as the average of at least two independent experiments, each performed in triplicate. Error bars represent the SEM.

### HOXA2 and HOXA3 select variants of the HOX/PBX motif

The observations above indicate that HOXA2 and HOXA3 select different genomic sites *in vivo*, while at a first glance, they recognize very similar DNA sequences. To investigate the determinants of HOX binding specificity, we focused on high confidence HOXA2 and HOXA3 peaks, which display the lowest overlap across the genome (Fig. 1E). *De novo* motif discovery identified enrichment of a HOX-PBX variant in HOXA3 top 250 peaks, which contains a C in the second variable position (i.e. TGAT**NC**AT) (Fig. 2A). We next counted the distribution of all permutations of the TGATNNAT motif in top HOXA2 and HOXA3 peaks and found the TGAT**TC**AT variant to be highly differentially enriched in HOXA3 peaks (Fig. 2B). This sequence, which is highly represented in HOXA3 top peaks (∼ 20%), is almost excluded from HOXA2 peaks (Fig. 2B). Supporting functional significance, HOXA3 peaks containing TGAT**TC**AT display increased acetylation levels (a mark of active enhancers) (Creyghton et al., 2010) in HOXA3-expressing tissues (Fig. 2C). In addition, while HOXA2 peaks display a very high representation of TGAT**GG**AT and TGAT**TG**AT, HOXA3 high confidence binding allows higher variability (four variants are counted > 20 times in HOXA3 peaks as opposed to only two variants in top HOXA2 peaks) (Fig. 2B). The highest differential enrichment of TGATNNAT variants is observed in top HOXA2 and HOXA3 peaks (Fig. 2-Supplemental Fig. 1A), which also display minimal overlap across the genome (Fig. 1E); this suggests that the ability to recognize different sequences plays a role in genomic site selections. Finally, the majority of HOXA3 (158/250) and HOXA2 (160/250) top peaks contain MEIS recognition motif, at a preferential distance of less than 20 nt from the TGATNNAT motif (Fig. 2-Supplemental Fig. 1B). The *Sulf2* locus exemplifies HOXA3 specific binding in PBA: it contains a single TGAT**TC**AT motif and displays high HOXA3 occupancy, but no detectable HOXA2 binding (Fig. 2DE). We used electrophoretic mobility shift assay (EMSA) to establish if HOXA3 preferentially recognizes the TGATTCAT sequence *in vitro*. We did not observe any HOXA2 or HOXA3 binding to the *Sulf2* probe (Fig. 2F). Incubation with PBX and MEIS resulted in a probe shift. Addition of HOXA3, but not HOXA2, resulted in the formation of a ternary complex, indicating that HOXA3 can bind this site in combination with PBX and MEIS, while HOXA2 cannot (Fig. 2F). In support of this conclusion, converting TGATT**C**AT to TGATT**G**AT (a single nucleotide substitution in the *Sulf2* probe), enables binding of HOXA2, in addition to HOXA3 (Fig. 2G). These results indicate that HOXA3 and HOXA2 have diverse binding preferences and uncover the existence of sites that are exclusively recognized by HOXA3.

### HOXA2 molecular control of BA2 identity

In contrast to HOXA3, which displays unique binding preferences for TGATTCAT, we did not detect HOX-PBX variants exclusively recognized by HOXA2. To investigate the mechanisms underlying HOXA2 control of BA2 identity, we examined HOXA2 binding events (top peaks) in the vicinity of well-established HOXA2 downstream targets. *Meis2* and *Zfp703* are associated with high levels of HOXA2 binding (Amin et al., 2015) (Fig. 3A and Fig. 3-Supplemental Fig. 1A) and are downregulated in *Hoxa2* null BA2 (Donaldson et al., 2012). In addition, consistent with *Meis2* and *Zfp703* expression being HOXA2-dependent, they are expressed at higher levels in BA2 than the HOX-less BA1 and the HOXA3-positive PBA (Fig. 3B). *Meis2* and *Zfp703* loci exhibit high HOXA2 and HOXA3 binding in their vicinity, suggesting their associated chromatin is largely accessible in both BA2 and PBA (Fig. 3A and Fig. 3-Supplemental Fig. 1A). We focused primarily on the *Meis2* enhancer, which is active in the main domains of HOXA2 expression, the hindbrain and BAs in zebrafish (Fig. 3C). When tested in a luciferase assay, the *Meis2* functional enhancer displays higher activity in the presence of HOXA2, in combination with MEIS and PBX, relative to HOXA3 (Fig. 3D). *Meis2* enhancer activity is strictly dependent on the integrity of its HOX-PBX site (Fig. 3D and Fig. 3F). Similar results were obtained with *Zfp703* putative enhancer, however in this case, HOXA2 and HOXA3 alone resulted in higher activation, presumably due to the presence of additional TAAT sites around the HOX/PBX motif (Fig. 3-Supplemental Fig. 1B). As for the *Meis2* enhancer, disruption of the HOX/PBX site nearly abolished activation (Fig. 3-Supplemental Fig. 1B). Finally, HOXD3, another HOX paralog group 3, also displayed a lower activating capacity than HOXA2 (Fig. 3-Supplemental Fig. 1C). In sum, HOXA2 is more efficient at activating both target regions, in the presence of PBX and MEIS. To understand if this reflects HOXA2 and HOXA3 different DNA binding properties, we generated HOX chimeric proteins by swapping HOXA2 and HOXA3 DNA-binding HDs. We found that providing HOXA2 with HOXA3 HD did not substantially change the ability of HOXA2 to activate transcription from the *Meis2* enhancer (Fig. 3E). Similarly, the ability of HOXA3 to transactivate the *Meis2* and *Zfp703* enhancers, alone or in complex with MEIS and PBX, was not improved by swapping HOXA3 HD with HOXA2 HD (Fig. 3E and Fig. 3-Supplemental Fig. 1B). As HOX TFs cooperate with MEIS and PBX to activate target enhancers and activation relies on the presence of an intact HOX/PBX motif, HOXA2 and HOXA3 diverse activation properties may depend on their respective abilities to interact with PBX and MEIS on DNA. On their own, HOXA2 and HOXA3 weakly bind the *Meis2* enhancer, but interact with PBX and MEIS to form a ternary protein complex on DNA (Fig. 3G-H). A larger fraction of MEIS-PBX complex is bound by HOXA2, while addition of HOXA3 result in a less robust supershift (Fig. 3GH). We observed the same binding patterns using HOX chimeras: swapping HOXA3-HD with HOXA2-HD did not improve the ability of HOXA3 to form a ternary complex with PBX and MEIS, and did not affect HOXA2 ability to bind DNA in complex with MEIS and PBX (Fig. 3I). Finally, altering the sequence of the HOX-PBX motif abolished formation of a HOX-MEIS-PBX complex on DNA (Fig. 3J). These results indicate that the differential ability of HOXA2 and HOXA3 to bind and activate transcription does not depend on HOX-DNA binary binding. Rather, it reflects differential abilities to form functional HOX-TALE complexes on DNA and is encoded by residues outside the HOXA2 and HOXA3 HD. In summary, while HOXA2 does not exclusively access its sites (HOXA3 can bind as well, Fig. 3A), HOXA2 binds more efficiently with TALE at these sites, leading to increased transcriptional activation. Consistently, shared high-confidence HOXA2 and HOXA3 binding events are largely associated with genes expressed at higher levels in the BA2 (Fig. 3K). Thus, at least in part, HOXA2 instructs the formation of a BA2 by raising the expression levels of HOX-regulated genes. Crucially, among these genes is *Meis2*, which encodes a critical component for BA2 identity (Amin et al., 2015).

### HOXA2 activity is decreased in PBA

The above results show that HOXA2 functions more efficiently with TALE relative to HOXA3. Given that HOXA2 is expressed in both the BA2 and in the PBA, why does HOXA2 not instruct a BA2-specific program in the PBA as well? More posterior *Hox* genes are typically able to repress the expression (and suppress the function) of more anterior genes, a process termed ‘posterior prevalence’ (Duboule, 2007). Indeed, *Hoxa2* highest expression is detected in the BA2, while *Hoxa2* is expressed at lower levels in *Hoxa3* main domain of expression, the BA3 (Fig. 4AB and Fig. 1B). To assess how changes in HOXA2 dose affect binding genome-wide, we compared HOXA2 binding in BA2 and in PBA. While HOXA2 binds similar locations in BA2 and PBA (Fig. 1H), HOXA2 binding levels are typically higher in BA2 (Fig. 4C, see also Fig. 3- Supplemental Fig. 1A). This is further confirmed by quantitative analysis of selected regions (Fig. 4D). Relative to BA2 cells, cells in the PBA display lower levels of HOXA2 and also express HOXA3 (Fig. 1B). We investigated the effect of decreasing HOXA2 levels and increasing HOXA3 levels on HOXA2 target enhancers. We found that co-expressing HOXA2 and HOXA3 reduced activation of HOXA2 target enhancers *in vitro* (Fig. 4E). In conclusion, a lower dose of HOXA2 decreases HOXA2 binding and activating abilities. This effect, combined with the lower efficiency of HOXA3 to activate HOXA2 targets, dampens HOXA2 transcriptional program in the PBA.

### HOX directly cooperates with MEIS

Our results indicate that HOX selectivity is displayed in concert with TALE. Generally, binding with TALE appears to be a dominant feature of HOX binding in the BAs. HOX peaks are enriched in HOX-PBX and MEIS motifs and similar to HOXA2 in BA2 (Amin et al., 2015), HOXA3 peaks overlap almost entirely with MEIS and PBX peaks in the same embryonic tissue at the same stage (Fig. 5A, Fig. 5-Supplemental Fig. 1A). We previously discovered that HOXA2 switches its transcriptional program by increasing binding of MEIS TFs to potentially lower-affinity sites across the genome (Amin et al., 2015). We investigated if HOXA3 can similarly increase MEIS binding levels. The fraction of MEIS peaks that overlaps HOXA3 binding displays higher FE in PBA, relative to the HOX-free BA1 (Fig. 5B). *Hoxa2* is also expressed in PBA, where it could be entirely responsible for the observed increase in MEIS binding. Therefore, to assess HOXA3 unique contribution to MEIS binding increase, we extracted HOXA3-specific binding. We found that MEIS peaks in PBA that overlap HOXA3 ‘exclusive’ peaks, display higher FE (relative to MEIS non-overlapping HOX), indicating that HOXA3 also increases binding of MEIS (Fig. 5C), similar to HOXA2 in BA2 (Amin et al., 2015) (FigS5). Reciprocally, co-occupancy with MEIS enhances HOXA3 binding (Fig. 5D). Both HOXA2 and HOXA3 interact with MEIS1 and MEIS2 (Fig. 5E), identifying direct cooperativity as the underlying mechanism. Direct cooperativity with MEIS appears to be a general operational principle of HOX TFs as, similar to HOXA2 and HOXA3, MEIS co-occupancy with HOXA1 and HOXA9 is associated with the highest MEIS binding levels in mouse embryonic stem cells (De Kumar et al., 2017) and bone marrow cells (Huang et al., 2012) respectively (Fig. 5-Supplemental Fig. 1B-D). In sum, HOX directly cooperate with TALE on chromatin. As HOXA2 and HOXA3 display sequence preferences and diverse binding affinities, HOX paralogs preferentially cooperate with distinct subsets of TALE binding events.

**Figure 5.**
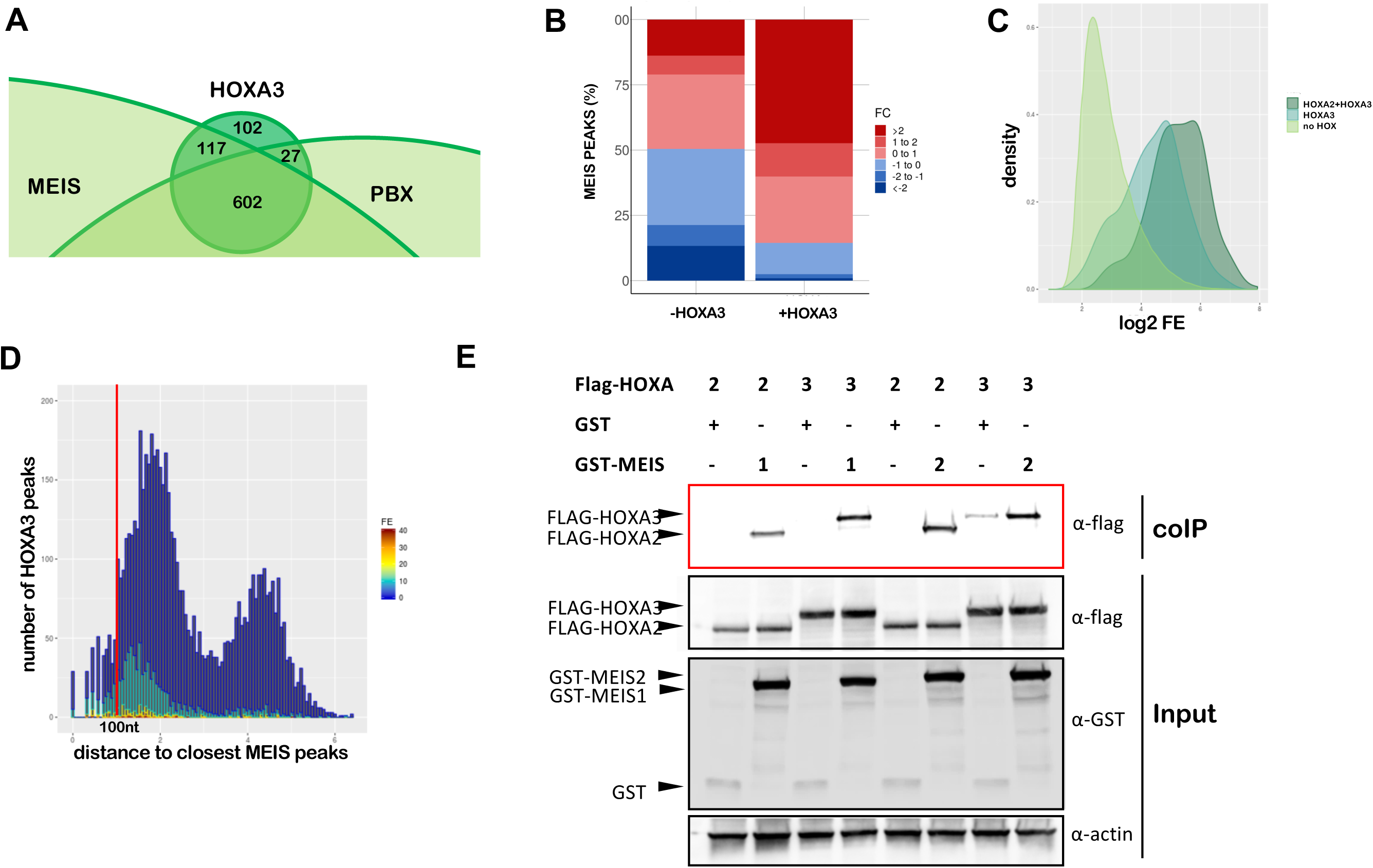
HOX directly cooperate with MEIS. A. Overlap of HOXA3 with MEIS and PBX peaks in the same tissue (PBA) and at the same embryonic stage (E11.5) (200nt summit regions, overlap at least 1nt). The proportional Venn diagram is cropped to focus on HOXA3 peaks. B. Barplots of fold change in MEIS binding levels in PBA versus BA1. Regions co-occupied by MEIS with HOXA3 in PBA generally display higher MEIS binding levels in PBA (HOX-positive) relative to the HOX-negative BA1. In contrast, MEIS binding not overlapping HOXA3 can be higher in BA1 or in PBA. Fold changes were calculated using EdgeR (see also Figure 6-figure Supplement 1). C. Kernel density plots of MEIS peaks relative to FE (PBA). MEIS binding is sorted into peaks not overlapping HOX (light green), MEIS peaks overlapping HOXA3 only (‘exclusive’ peaks, i.e. not overlapping HOXA2 in PBA, darker green) and MEIS peaks overlapping HOXA2 and HOXA3 (darkest green). D. Distance of HOXA3 peaks relative to MEIS peaks (PBA). HOXA3 peaks are binned according to their log_10_ distance to the nearest MEIS peak and labelled according to FE (high FE, dark red bars; low FE, dark blue bars). E. Co-immunoprecipitation assays. HEK293T cells were co-transfected with expression vectors for FLAG-tagged HOXA2 or HOXA3 and GST-tagged MEIS1, GST-tagged MEIS2 or GST alone. Protein interactions were assayed by co-immunoprecipitation on glutathione beads directed toward the GST tag and eluted proteins analysed by western blotting to detect the presence of HOXA2-FLAG or HOXA3-FLAG (red box, Co-IP). Cell lysates were analysed by western blotting prior to co-immuno precipitation to detect protein expression (input).

### MEIS ‘ubiquitous’ binding is converted into tissue-specific enhancer activity

MEIS TFs bind broadly and to largely overlapping locations across different BAs (Fig. 6A) (Amin et al., 2015), and only a small fraction of TALE-bound regions is occupied by HOX (Fig. 5-Supplemental Fig. 1A). HOX-MEIS cooperativity predicts that the fraction of high MEIS peaks in HOX-positive areas (BA2 and PBA), should be enriched in HOX motifs. We systematically extracted differential MEIS binding across the BAs (Fig. 6-Supplemental Fig.1) and found, using convolutional neural network (CNN) models, that differential classification of MEIS binding is sufficient to uncover HOX motif features (Phuycharoen et al., 2019); specifically, the fraction of MEIS peaks higher in BA2 and in PBA (= lower BA1) is highly enriched in sequence features matching HOX-PBX motif (Fig. 6B). Interestingly, the same CNN models identify enrichment of other TF recognition motifs in differential MEIS binding (Fig. 6B). These signature motifs reflect a differential distribution of TFs across the BAs (Fig. 6C). Moreover, CNN models detect established TF interactions (Jolma et al., 2015), as well as TF co-occupancy detected *in vivo* (Losa et al., 2017). Namely, GATA recognition motifs are enriched in higher MEIS binding in PBA, and GATA TFs are exclusively expressed in PBA (Fig. 6C), where GATA6 and MEIS bind overlapping locations. These observations suggest that other tissue-specific TFs, in addition to HOX, can affect MEIS binding to chromatin. Next, we globally quantified changes in enhancer activity across the BAs to assess the function of MEIS differential binding. Consistent with MEIS positive effects on transcription (Choe et al., 2009), regions occupied by HOXA2 in BA2, or HOXA3 in PBA, display higher enhancer activity when associated with increased MEIS binding levels in the same tissue (Fig. 6D). More generally, higher MEIS binding levels in a tissue are highly predictive of increased enhancer activity in the same tissue (Fig. 6E), an effect only partly explained by HOX-MEIS cooperativity (Fig. 6-Supplemental Fig. 2AB). Finally, supporting the concept that MEIS ubiquitous binding (Fig. 6A) is transformed into BA-specific enhancer activity, top MEIS binding is BA-specific and associated with distinct biological processes (Fig. 6FG and Fig. 6-Supplemental Fig. 2C). De novo motif discovery on HOXA3-and HOXA2-specific peaks identifies enrichment of distinctive sequence features of MEIS differential binding in PBA and BA2, NKX (HD) and FOX (Forkhead) motifs and basic helix-loop-helix (bHLH) recognition sites respectively (Fig. 6H), suggesting that HOX and tissue-specific TFs may collaborate in binding with TALE. We focused on FOX TFs, because *Fox* genes are typically expressed at higher levels in PBA than BA2 (Fig. 6C). Consistent with the three factors cooperating on chromatin, HOX and FOX recognition sites co-occur in the same differential MEIS peaks (Fig. 6-Supplemental Fig. 2DE). Moreover, FOXC1 binding in the BA (Amin et al., 2015) partly overlaps with HOXA2 and HOXA3 binding (Fig. 6-Supplemental Fig. 2F). FOXC1, HOX and MEIS/PBX synergize to increase transcriptional activation driven by the *Sfrp2* distal region (co-occupied by HOX and FOXC1) (Fig. 6I). Interestingly, the presence of FOXC1 is sufficient to enhance MEIS/HOX transcriptional activation of *Sfrp2* enhancer, suggesting that cooperation between these TFs could partly compensate for lack of PBX (Fig. 6I). While FOXC1 display similar cooperativity with TALE and HOXA2 or HOXA3 *in vitro*, the higher levels of FOX TFs in the PBA, relative to BA2, predict FOX TFs to have stronger effects on HOXA3 and MEIS binding in PBA; this expectation is supported by the enrichment of FOX motifs in HOXA3 and MEIS differential binding in PBA, but not HOXA2 and MEIS differential binding in BA2 (Fig. 6BH). Indeed, in silico mutagenesis predicts mutations in FOX TF recognition sites to affect binding of both HOXA3 and MEIS in PBA, but not HOXA2 and MEIS in BA2 (Fig. 6J, Fig. 6-Supplemental Fig. 2G). In contrast, mutagenesis of GATA motifs (enriched in MEIS differential peaks, but not in HOX peaks) does not appear to affect HOX-MEIS binding (Fig. 6J). These results identify (direct or indirect) cooperativity with tissue-specific TFs as an additional mechanism for HOX selectivity. We propose that HOX and tissue-specific TFs (alone and in combination) increase TALE TF binding affinity and residence time at selected locations, identified using their sequence recognition motifs. Increasing MEIS residence time on chromatin has a positive effect on enhancer activity and results in BA-specific transcriptional outputs. Thus, TALE TFs function as a hub which integrates different signals instructing BA morphogenesis.

**Figure 6.**
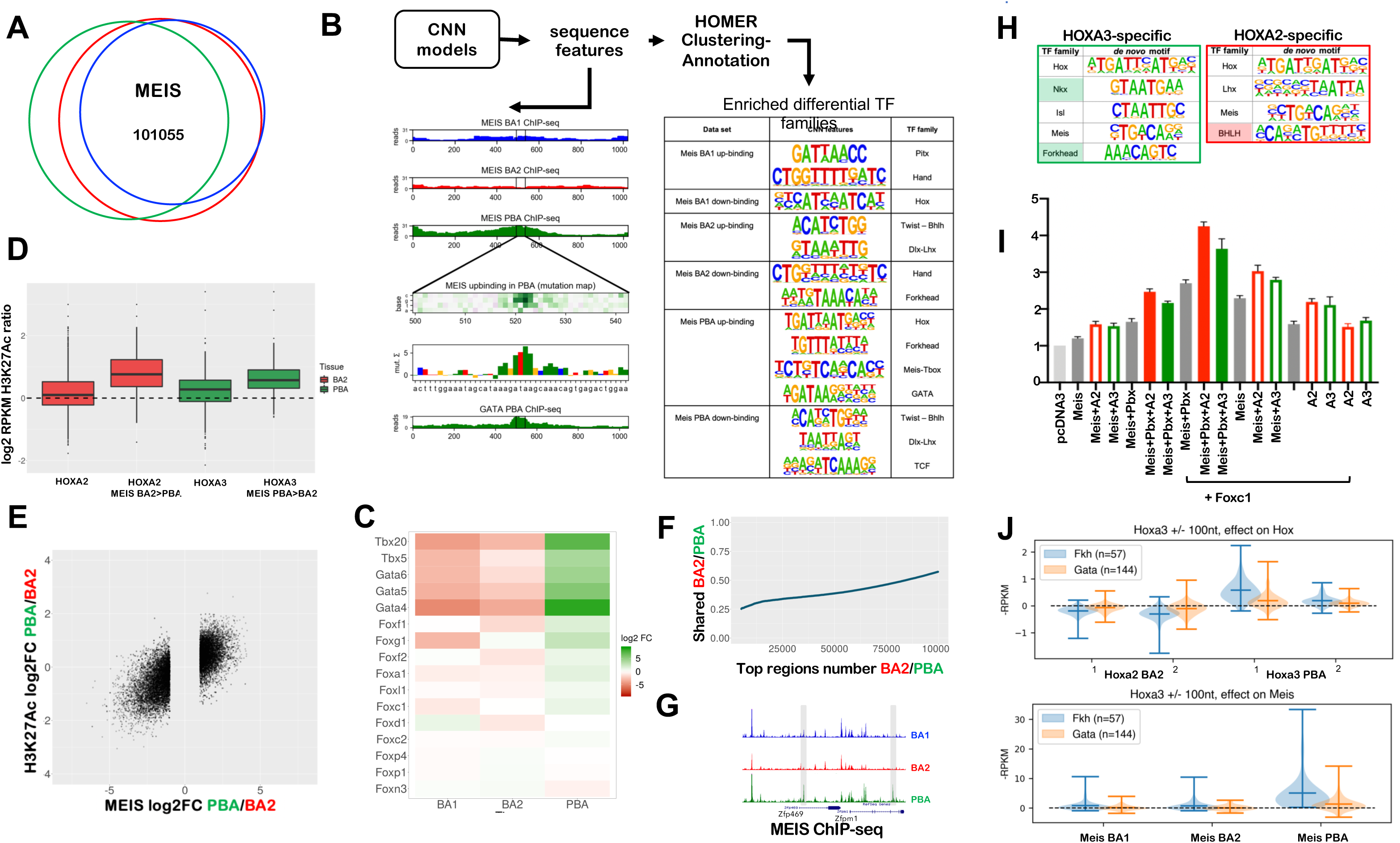
A. Proportional Venn diagram shows highly overlapping binding of MEIS in BA1, BA2 and PBA. Out of 215830 MEIS peaks, 101055 are in common between the three tissues; MEIS peaks were combined and re-centered using DiffBind. B. CNN models of MEIS differential peaks uncover enrichment of tissue-specific sequence motifs as described in (Phuycharoen et al., 2019). MEIS binding was classified in six categories (i.e. peaks with higher/lower binding in BA1, BA2, PBA). CNN analysis identifies tissue-specific sequence features in each class of MEIS peaks. Predicted GATA binding in a MEIS PBA up-binding region is visualised as in the example (a feature matching GATA TF recognition motif on chr5:104257972-104258015 is shown) and annotated using HOMER. The GATA6 ChIP-seq verifies this prediction. HOMER was used to cluster and annotate tissue-specific sequence features; differentially enriched features are matched to TF families with known tissue-specificity (see also Fig. 6C). C. Heatmap of the expression of selected TF families, corresponding to cognate recognition motifs identified in MEIS PBA-up, in E11.5 mouse BA2 and PBA. Members of the GATA and TBX families, and the majority of expressed Forkhead TFs are enriched in PBA relative to BA2. Only TFs with expression values > 10 cpm in at least one tissue are shown. D. Boxplots of the ratio of H3K27ac (log_2_RPKM) in BA2 and PBA for all HOX peaks and for HOX peaks overlapping MEIS differential binding higher in BA2 (HOXA2 peaks) and higher in PBA (HOXA3 peaks). HOX binding generally increases H3K27Ac; peaks associated with increased MEIS binding display a higher increment of H3K27Ac in the same tissue. E. Correlation plot of differential MEIS binding and differential acetylation (enhancer activity) at intergenic regions (PBA versus BA2). Each point corresponds to a region with MEIS log_2_ fold change >1 (FC>2); the corresponding H3K27ac value is plotted. Changes in MEIS binding levels are positively correlated with increased enhancer activity in the same tissue (correlation = 0.73). F. Different top MEIS peaks are observed in different BAs. The ratio of MEIS peaks, which are common to BA2 and PBA, increases as FE decreases. G. UCSC tracks illustrates MEIS increased binding at the *Zfp496* and *Zfpm*1 loci. Instances of common MEIS peaks higher in one tissue (PBA) are shaded. H. HOMER de novo motif discovery in HOXA3-specific and HOXA2-specific peaks. HOXA3-specific are HOXA3 peaks excluding peaks overlapping with HOXA2 BA2; similarly, HOXA2-specific are HOXA2 peaks excluding peaks overlapping with HOXA3 PBA. HOMER identifies enrichment of the same motifs enriched in BA-specific MEIS differential binding, Forkhead motif in HOXA3-specific (shaded in green) and BHLH motif in HOXA2-specific subsets (shaded in red). Variations of HD recognition motifs potentially recognized by HOX and attributed by HOMER to PBA-specific TFs NKX and ISL1 in PBA and LHX/DLX in BA2 are also enriched. I. Luciferase activity driven by *Sfrp2* enhancer co-transfected with *Meis*, and *Meis* and *Pb*x with and without *Hoxa2* (red empty bars), *Hoxa3* (green empty bars) and *Foxc1* (grey) in 3T3 cells. Adding *Foxc1* to *Hoxa2* or *Hoxa3* with *Meis2* and *Pbx1a* results in the highest activation. J. In silico knockout of Forkhead and GATA motifs is used to predict the effects on HOX and MEIS binding. CNN MEIS PBA ‘up-binding’ features (Fig. 6B) were annotated as HOX, GATA, and Forkhead (see methods). Co-occurring HOX-Forkhead motifs (distance between 1 nt to 100 nt) were selected for in silico mutagenesis. Forkhead mutagenesis results in a significant drop in HOXA3 binding in PBA, but shows no average significant effect on HOXA2 in BA2. Similarly, Forkhead mutagenesis significantly decreases Meis PBA binding across most tested sites. In comparison, much weaker effects are predicted on BA1 and BA2 MEIS differential binding. As a negative control, the same procedure was applied to co-occurring HOX-GATA motifs. GATA motif mutagenesis does not show significant average effects on HOX, or MEIS in HOX-bound regions.

## Discussion

HOX TFs contain a HD, which display highly similar sequence recognition properties and is shared by hundreds of TFs, yet they instruct diverse, segment-specific transcriptional programs along the antero-posterior axis of all bilaterian animals. By profiling HOXA2 and HOXA3 binding in their physiological domains, we identify three main determinants of HOX-selective binding across the genome: 1) recognition of unique variants of the HOX-PBX motif; 2) differential affinity at ‘shared’ HOX-PBX motifs and; 3) presence of additional tissue-specific, non-TALE, TFs. These mechanisms (with the possible exception of the first) are expected to generate quantitative (rather than qualitative, i.e. binding/no binding) differences in the relative levels of HOX/TALE occupancy on commonly bound regions. Such quantitative changes are a feature of continuous networks (Biggin, 2011), in which TFs bind a continuum of functional and non-functional sites and regulatory specificities derive from quantitative differences in DNA occupancy patterns.

HOX paralog-selective binding occurs in cooperation with TALE. The high degree of HOX and TALE interaction flexibility, mediated by paralog-specific protein signatures, has been proposed to generate paralog-specific functions of HOX TFs (Dard et al., 2018). Here, by defining the *in vivo* repertoire of HOX occupied sites, we identify DNA sequence as an additional determinant of HOX-TALE functional specificity *in vivo*. This finding is consistent with the mechanism of latent specificity described for *Drosophila* Hox/Exd (PBX) interaction (Slattery et al., 2011) and *in vitro* observations that HOX TFs bind longer, more specific sequence motifs in the presence of TALE. However, the effects of TALE on HOX binding *in vivo* go beyond the refinement of HOX binding sites as, at least in the BA context, binding with TALE appears to be a requirement for loading HOX on chromatin. Our observations indicate that HOXA2-A3 overwhelmingly recognize genomic sites that are enriched in HOX-PBX motifs and are also occupied by TALE TFs *in vivo*. Therefore, TALE provides a platform for HOX to bind; selectivity enables HOX paralogs to preferentially bind different subsets of this common platform. In agreement with our finding that BA-specific chromatin states do not seem to play a role in HOX target site selection, TALE platform is largely similar across BA1-2-PBA.

What is the functional significance of HOX-TALE interaction on chromatin and how does it contribute to paralog-specific transcriptional programs? Many examples from animal development indicate that transcriptional regulation is mediated by distinct combinations of TFs. TALE TFs operate as a hub, which assists combinatorial assembly of TF complexes. TALE platform expands HOX functional interface and enables HOX to function in concert with other TFs, bypassing the need of direct protein-protein interaction. In doing so, it integrates positional signals (encoded by HOX) and local inputs (provided by cell type-/tissue-specific TFs) into defined transcriptional outputs. While it is possible that MEIS and PBX facilitate access of diverse TFs to relatively inaccessible chromatin, MEIS TFs differ from conventional pioneer TFs, which function to open chromatin regions but are not directly involved in enhancer activation (Cirillo et al., 1998; Jacobs et al., 2018). Remarkably, independently of the type of TF involved (HOX or other tissue-specific TFs), positive changes in MEIS binding result in a functional effect, i.e. increased enhancer activity. High instances of MEIS binding are typically tissue-specific and highly correlated with enhancer activity. In fact, differential MEIS binding in a specific BAs is generally a very good predictor for matching changes in enhancer activity in the same tissue. Based on our observations and the well-established role of MEIS in transcriptional activation (Choe et al., 2009; Hau et al., 2017; Hyman-Walsh et al., 2010), we propose a model of transcriptional activation, where TALE (MEIS) TFs function as a broad or general activators and HOX paralog selectivity is mainly directed at harnessing TALE functional activity at selected locations. Using their recognition motifs, HOX and/or tissue-specific TFs select specific MEIS binding locations, where they stabilize MEIS binding to generate precise functional outputs, or patterns of enhancer activation (Fig. 7). Interestingly, MEIS2 interacts with PARP1 (Hau et al., 2017), a large enzyme capable of triggering phase condensation (Altmeyer et al., 2015). Increasing MEIS residence time (as a result of the cooperation with HOX and other TFs) may favour PARP1 recruitment at selected loci and, in turn, generate the liquid-liquid phase transitions observed to promote gene activation (Boija et al., 2018; Hnisz et al., 2017).

**Figure 7.**
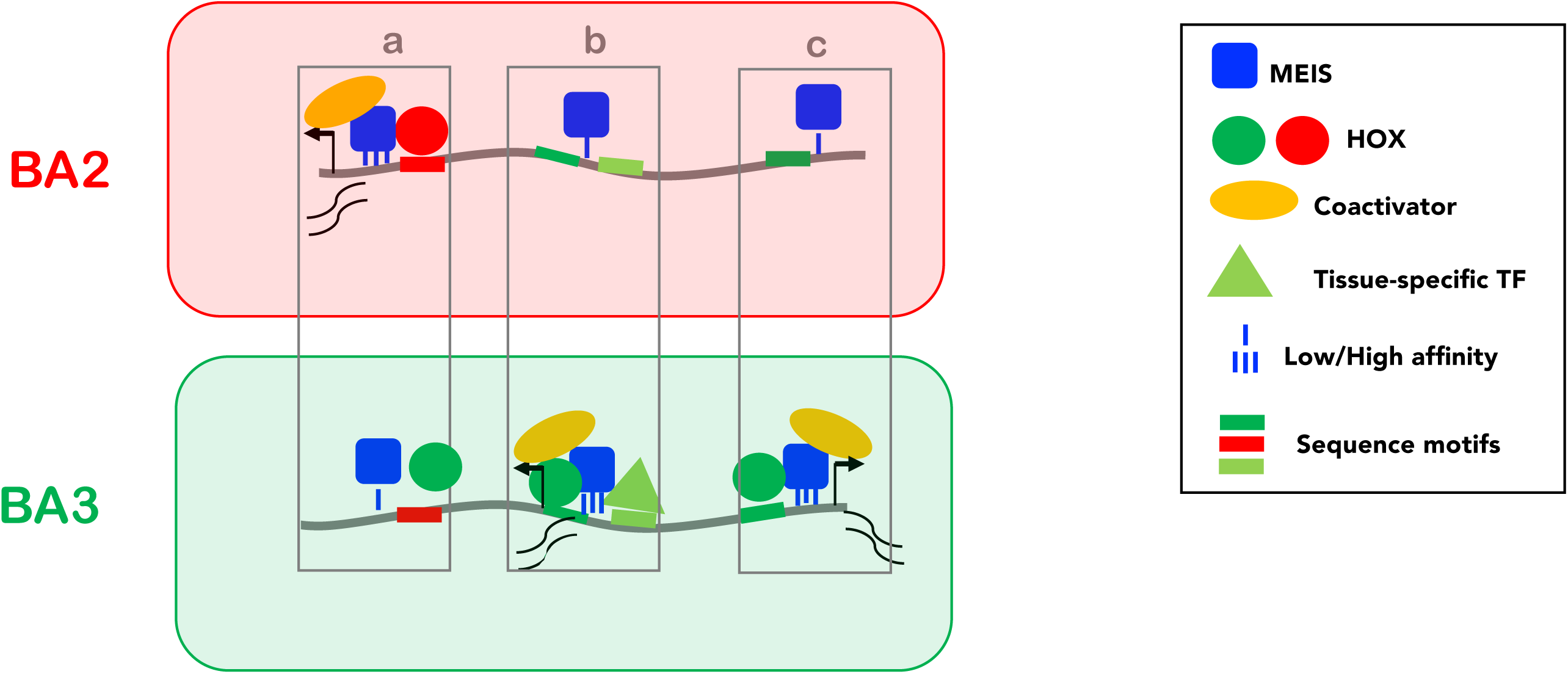
Model. Low affinity, widespread binding of MEIS (blue square) defines a large subset of accessible chromatin (grey line) for activation (PBX is not shown as PBX and MEIS binding almost entirely overlaps). Direct cooperativity with HOX (A2 and A3, red and green circles respectively) and/or indirect cooperativity with tissue-specific TFs (triangle) increase MEIS binding affinity and residence time; prolonged residence time of MEIS at enhancers promotes recruitment of general co-activators (yellow) and activation of transcription. HOX paralogs preferentially bind different subsets of MEIS occupied regions, resulting in differential transcription. Three examples of BA-specific transcription are shown. In **a**, the red site is bound with higher affinity by HOXA2 than HOXA3, resulting in the formation of a more stable HOX-TALE complex on DNA and a (higher) transcriptional output in BA2. Conversely, in **c**, the green site is only recognized by HOXA3, leading to high affinity MEIS binding only in PBA, and to PBA-specific transcription. In **b**, the effect of HOXA3 is potentiated by a PBA-specific TF binding in the vicinity. Co-binding with tissue-specific TFs may positively contribute to HOX-MEIS cooperativity by competing with nucleosome for DNA binding, especially at HOX and/or MEIS low affinity sites. These mechanisms result in BA-specific transcription.

Because high instances of MEIS binding are typically associated with combinatorial TF binding, a precise identification of the critical steps for enhancer activation, and their sequential order, remains problematic. For similar reasons, MEIS and PBX shared genomic occupancy complicates dissecting their respective contributions to enhancer binding and activation. In addition to TALE, numerous other TFs are broadly, if not ubiquitously expressed during development, yet their inactivation results in tissue-specific phenotypes. It is tempting to speculate that similar principles of TF functional connectivity could explain other transcriptional networks, i.e. that cell type-tissue-specific regulators harness the activation abilities of broadly expressed TFs to generate cell type -specific gene expression programs.

## Supporting information

Supplemental Figures with legends

## Acknowledgements

We thank Ian Donaldson, Andy Hayes and the other members of the Genomic Technologies, Bioinformatics and Biological Services Core Facilities at the University of Manchester. We also thank Rene Reszohazy for sharing plasmids and Samir Merabet for helpful suggestions. This work was supported by MRC grant MR/L009986/1 to NB, BBSRC grant BB/N00907X/1 to NB, MR and KM and a BBSRC studentship to ML. FL and CS were supported by NIH grant NS038183 to CS.

## Material and methods

### Animal experiments

CD1 mice were time-mated to obtain BA2 or PBA from E115 embryos. Mouse experiments were carried out under ASPA 1986. Wild type zebrafish were raised in the University of Massachusetts Medical Center Zebrafish Facility. Embryos and adult zebrafish were maintained under standard laboratory conditions. Enhancers were amplified from mouse genomic DNA using the primers (listed in S), cloned into pCR8/GW/TOPO vector (Life Technologies) and recombined using the Gateway system (Life Technologies) to an enhancer test vector that includes a strong midbrain enhancer (Minitol2-GwB-zgata2-GFP-48, a kind gift from JL Skarmeta) as an internal control. Fertilized zebrafish embryos were collected from natural spawnings. Plasmid DNA was injected into the cytoplasm of one-cell stage embryos. Injected embryos were visualized intermittently by fluorescence microscopy up to 48 hr post fertilization to identify transgenic carriers. These were raised to adulthood, outcrossed to wildtype fish and the resulting F1 embryos were scored for GFP expression in order to generate stable transgenic lines.

### Next-generation sequencing data and downstream analyses

ChIP-seq was performed as described (Losa et al., 2017) using rabbit polyclonal antibodies targeting HOXA3 (non-conserved N-terminal amino acids 24 to 180), HOXA2 (Kutejova et al., 2008), PBX1-2-3-4 (sc-25411X, Santa Cruz) and rabbit IgG (Millipore). DNA was recovered from two independent ChIP-seq experiments and purified using DiaPure columns (Diagenode). Enrichment was validated by SYBR green quantitative PCR (qPCR) using primers listed in Table S1. DNA libraries were constructed using the MicroPlex Library Preparation Kit v2 (Diagenode) and sequenced with the Illumina next generation sequencing platform. ChIP-seq experiments were analysed using Trimmomatic for trimming (Bolger et al., 2014), Bowtie2 for aligning to the mouse genome (mm9) (Langmead and Salzberg, 2012), samtools (Li et al., 2009) to remove the aligned reads with a mapping quality Q30 and MACS2 for peak calling (Zhang et al., 2008) with default narrow peak calling setting for TFs and broad peak calling setting for histone modification marks. ‘*findMotifGenome’* module of the HOMER package was used to detect *de novo* motif in 200nt summit regions (Heinz et al., 2010). Venn diagrams were generated using 200nt peak summits with an overlap of at least 1nt. GREAT standard association rule settings (McLean et al., 2010) was used to associate ChIP-seq peaks with genes and uncover events controlled by TF binding. DiffBind (Ross-Innes et al., 2012) was used to re-center MEIS and H3K27ac peaks across BA1, BA2 and PBA (Figure 6_supplemental Fig. 1) and calculate RPKM values and raw counts in the re-centered regions. edgeR generalized linear model (GLM) method with TMM normalization (Robinson et al., 2010) was used to select differential peaks and calculate fold change in MEIS binding and H3K27ac across BAs used to generate boxplots and scatterplots. The best H3K27Ac replicate [highest FRiPs (fraction of reads in peaks)] RPKM values was used to produce boxplots. Gene expression CPM values and differential gene expression at E10.5 and E11.5 were derived from (Amin et al., 2015; Losa et al., 2017). ggplot2 package (Wickham, 2016) was used to generate CPM values heatmap. GALAXY (Geocks et al 2010), Bioconductor GenomicRanges package (Lawrence et al., 2013), and Bioconductor ChIPpeakAnno package (https://www.bioconductor.org/packages//2.10/bioc/html/ChIPpeakAnno.html) were used to intersect, modify and visualize genomic coordinates. Bioconductor Biostring (Pagès H, 2019) was used to locate fixed motif sequences in the binding regions. Distance between HOX and MEIS binding regions was calculated using GenomicRanges package and plotted with ggplot2. The Kernel density distribution of MEIS fold enrichment in HOX binding regions vs non-HOX binding regions were calculated by R kernel density distribution estimation (R core team 2013) and plotted with ggplot2.

All RNA-seq and ChIP-seq datasets are available on the ArrayExpress with accession numbers: E-MTAB-7963, E-MTAB-7966, E-MTAB-7766, E-MTAB-7767, E-MTAB-5394, E-MTAB-5407, E-MTAB-5536, E-MTAB-2696.

### Convolutional neural network models and in silico mutagenesis

MEIS differential sequence features are detected by recently published differential convolutional neural network (CNN) structure (Phuycharoen et al., 2019). For in silico binding site knockout we trained a convolutional neural network (CNN) model for multitask regression of MEIS and HOX RPKM binding level. The CNN was trained by transfer learning, using convolution parameters from a previously published 1-convolutional layer MEIS RPKM model (Phuycharoen et al., 2019). Convolutional filters were transferred to a new model, which was then trained on a subset of MEIS regions also bound by HOX, to simultaneously predict log2RPKM values in 2 replicates of Hoxa2 in BA2, 2 from Hoxa3 in PBA, and one replicate of MEIS in BA1, BA2 and PBA. The training data consisted of 6795 regions of 600nt with HOX binding predicted by MACS2 in any tissue. The regression model was subsequently used to predict the change in RPKM values after binding site erasure. For simulated genomic knockout, a 25nt site containing each feature was replaced by random di-nucleotides from the remaining part of the region and RPKM levels were predicted. Random replacement was repeated 100 times for each feature, averaging the predicted RPKM change. To select candidate features for erasure, MEIS PBA up-binding features were first obtained from the previously published 3-task parallel model and subsequently filtered. Sites of HOXA3 and GATA were required to contain consensus motif “TGATNNAT” and “WGATAA” respectively, with no mismatch allowed. Forkhead sites were selected based on long distinct k-mers, derived from KSM motif representation method (Guo et al., 2018), namely exact matches to any of the following sequences: “AAAATAAACA”, “AAAAATAAAC”, “AATAAATCAA”, “ATNAATCAACA”, “AAATAAACAC”, “ATAAATCAAC”,”GAAAATAAAC”, “CAAAATAAAC”, “AAAATAAACT”, “AAATAAACAA”. These candidate sites were identified within a +/-250nt window centred on HOXA3 and GATA6 ChIP-seq peak summits, FE of replicates was combined with edgeR (Robinson et al. 2010) and a Poisson test was performed as in MACS2 using false discovery rate (FDR) cutoff = 0.05. Only Forkhead and GATA motifs that did not contain internal matches to HOX-PBX motif were selected. Subsets of GATA and Forkhead sites located within +/-100nt from a HOX-PBX sites were selected for mutagenesis.

### Elecrophoretic mobility shift assays

Probes were made from primers with 5’ ATO700, and purified with QIAGEN PCR purification kit (Qiagen). Proteins were generated using TnT® Quick Coupled Transcription/Translation System (Promega) and the following plasmids: pcDNA3-Hoxa2, pcDNA3-Hoxa3, pcDNA3-Meis2, containing mouse coding sequences for *Hoxa2, Hoxa3* and *Meis2* (isoform 1), cloned into pcDNA3 (Invitrogen); pcDNA3-PBX1a is a gift from Francesco Blasi. Reactions (4% Ficoll, 20mM HEPES, 37.5mM KCl, 1mM DTT, 0.1mM EDTA, 2ug Poly dI.dC, 16ng probe, and 2ul of TNT extracts in total volume of 10ul) were mixed by gentle flicking, and incubated at room temperature for 12 minutes before being run on 3% / 4% acrylamide gel at 70V in 0.5X TBE.

### Luciferase assay

*Meis2* and *Zfp703* enhancers were amplified from mouse genomic DNA using primers listed in Table S1 cloned into pCR8/GW/TOPO vector (Life Technologies) and recombined using the Gateway system (Life Technologies) into pGL4.23-GW (a gift from Jorge Ferrer; Addgene plasmid # 60323; http://n2t.net/addgene:60323; RRID:Addgene_60323). Enhancers were co-transfected with pcDNA3, pcDNA3-Hoxa2, pcDNA3-Hoxa3, pcDNA3-Meis2, pcDNA3-PBX1a (described above) and pcDNA3-Hoxd3 generated by GenScript. NIH3T3 cells were grown in DMEM (D6429) supplemented with 10% FBS and 5% penicillin/streptomycin, and seeded in 24-well plates at 100,000 cells/ml. Cells were transfected with GeneJuice Transfection Reagent (Novagen), using 250ng luciferase plasmid and 300ng pcDNA3 plasmids per well. Cells were harvested 24 hours after transfection and luciferase measured using Luciferase Assay System and the GloMax Multi-Detection System (Promega).

### Antibody validation

Gateway® entry vectors for mouse *Hoxb1* and *Hoxb2* (Bridoux & al. 2015 PubMed PMID: 26303204), human *HOXA3* and *HOXC4* (http://horfdb.dfci.harvard.edu/hv7/) were used to generate mammalian expression vectors for FLAG-HOX (v1899 destination vector) using the gateway technology (Barrios-Rodiles et al., 2005). Gateway® expression vectors for pExpFLAG-Hoxa1 and pExpFLAG-Hoxa2 are described in (Bergiers et al., 2013; Lambert et al., 2012). HEK293 cells were grown at 37°C, in a humidified atmosphere with 5% CO2 in DMEM (D6429) supplemented with 10% FBS, 5% penicillin/streptomycin, and 5% L-glutamine. Cells were seeded in 6-well plates at 400,000 cells/well and transfected 24 hours after plating using 1µg of HOX plasmid constructs and Fugene6 (Promega) according to the manufacturer’s instructions. Proteins were collected 48 hours after transfection, boiled in Laemmli buffer, run on SDS-page and visualized using anti-FLAG (M2) (#F1804, Sigma), HRP-conjugated anti-β-ACTIN (#A3854, Sigma) and anti-Hoxa3 antibody (1:2000) and HRP-HRP-conjugated secondary antibodies.

### Co-immunoprecipitation experiments

Coding sequences for MEIS1b and MEIS2.1 were cloned in pEnt plasmids, confirmed by DNA sequencing and used to generate pExp mammalian expression vectors for GST-tagged proteins with the pDest-GST N-terminal destination vector using the gateway technology (Rual et al., 2005). HEK293 cells were transfected as above, using 500ng each of FLAG/GST constructs per well. Proteins were collected 48 hours after transfection and co-precipitation performed as described in (Bridoux et al., 2015).

## References

Altmeyer, M., Neelsen, K.J., Teloni, F., Pozdnyakova, I., Pellegrino, S., Grofte, M., Rask, M.D., Streicher, W., Jungmichel, S., Nielsen, M.L., et al. (2015). Liquid demixing of intrinsically disordered proteins is seeded by poly(ADP-ribose). Nat Commun 6, 8088.

Amin, S., Donaldson, I.J., Zannino, D.A., Hensman, J., Rattray, M., Losa, M., Spitz, F., Ladam, F., Sagerstrom, C., and Bobola, N. (2015). Hoxa2 Selectively Enhances Meis Binding to Change a Branchial Arch Ground State. Dev Cell 32, 265–277.

Barrios-Rodiles, M., Brown, K.R., Ozdamar, B., Bose, R., Liu, Z., Donovan, R.S., Shinjo, F., Liu, Y., Dembowy, J., Taylor, I.W., et al. (2005). High-throughput mapping of a dynamic signaling network in mammalian cells. Science 307, 1621–1625.

Bergiers, I., Bridoux, L., Nguyen, N., Twizere, J.C., and Rezsohazy, R. (2013). The homeodomain transcription factor Hoxa2 interacts with and promotes the proteasomal degradation of the E3 ubiquitin protein ligase RCHY1. Plos One 8, e80387.

Biggin, M.D. (2011). Animal transcription networks as highly connected, quantitative continua. Dev Cell 21, 611–626.

Bobola, N., and Merabet, S. (2017). Homeodomain proteins in action: similar DNA binding preferences, highly variable connectivity. Curr Opin Genet Dev 43, 1–8.

Boija, A., Klein, I.A., Sabari, B.R., Dall’Agnese, A., Coffey, E.L., Zamudio, A.V., Li, C.H., Shrinivas, K., Manteiga, J.C., Hannett, N.M., et al. (2018). Transcription Factors Activate Genes through the Phase-Separation Capacity of Their Activation Domains. Cell 175, 1842–1855 e1816.

Bolger, A.M., Lohse, M., and Usadel, B. (2014). Trimmomatic: a flexible trimmer for Illumina sequence data. Bioinformatics 30, 2114–2120.

Bridoux, L., Bergiers, I., Draime, A., Halbout, M., Deneyer, N., Twizere, J.C., and Rezsohazy, R. (2015). KPC2 relocalizes HOXA2 to the cytoplasm and decreases its transcriptional activity. Biochim Biophys Acta 1849, 1298–1311.

Burglin, T.R., and Affolter, M. (2016). Homeodomain proteins: an update. Chromosoma 125, 497–521.

Choe, S.K., Lu, P., Nakamura, M., Lee, J., and Sagerstrom, C.G. (2009). Meis cofactors control HDAC and CBP accessibility at Hox-regulated promoters during zebrafish embryogenesis. Dev Cell 17, 561–567.

Cirillo, L.A., McPherson, C.E., Bossard, P., Stevens, K., Cherian, S., Shim, E.Y., Clark, K.L., Burley, S.K., and Zaret, K.S. (1998). Binding of the winged-helix transcription factor HNF3 to a linker histone site on the nucleosome. Embo J 17, 244–254.

Creyghton, M.P., Cheng, A.W., Welstead, G.G., Kooistra, T., Carey, B.W., Steine, E.J., Hanna, J., Lodato, M.A., Frampton, G.M., Sharp, P.A., et al. (2010). Histone H3K27ac separates active from poised enhancers and predicts developmental state. Proc Natl Acad Sci U S A 107, 21931–21936.

Cumberworth, A., Lamour, G., Babu, M.M., and Gsponer, J. (2013). Promiscuity as a functional trait: intrinsically disordered regions as central players of interactomes. Biochem J 454, 361–369.

Dard, A., Reboulet, J., Jia, Y., Bleicher, F., Duffraisse, M., Vanaker, J.M., Forcet, C., and Merabet, S. (2018). Human HOX Proteins Use Diverse and Context-Dependent Motifs to Interact with TALE Class Cofactors. Cell Rep 22, 3058–3071.

De Kumar, B., Parker, H.J., Paulson, A., Parrish, M.E., Pushel, I., Singh, N.P., Zhang, Y., Slaughter, B.D., Unruh, J.R., Florens, L., et al. (2017). HOXA1 and TALE proteins display cross-regulatory interactions and form a combinatorial binding code on HOXA1 targets. Genome Res 27, 1501–1512.

Donaldson, I.J., Amin, S., Hensman, J.J., Kutejova, E., Rattray, M., Lawrence, N., Hayes, A., Ward, C.M., and Bobola, N. (2012). Genome-wide occupancy links Hoxa2 to Wnt-beta-catenin signaling in mouse embryonic development. Nucleic Acids Res 40, 3990–4001.

Duboule, D. (2007). The rise and fall of Hox gene clusters. Development 134, 2549–2560.

Farley, E.K., Olson, K.M., Zhang, W., Brandt, A.J., Rokhsar, D.S., and Levine, M.S. (2015). Suboptimization of developmental enhancers. Science 350, 325–328.

Gendron-Maguire, M., Mallo, M., Zhang, M., and Gridley, T. (1993). Hoxa-2 mutant mice exhibit homeotic transformation of skeletal elements derived from cranial neural crest. Cell 75, 1317–1331.

Guo, Y., Tian, K., Zeng, H., Guo, X., and Gifford, D.K. (2018). A novel k-mer set memory (KSM) motif representation improves regulatory variant prediction. Genome Res 28, 891–900.

Hau, A.C., Grebbin, B.M., Agoston, Z., Anders-Maurer, M., Muller, T., Gross, A., Kolb, J., Langer, J.D., Doring, C., and Schulte, D. (2017). MEIS homeodomain proteins facilitate PARP1/ARTD1-mediated eviction of histone H1. J Cell Biol 216, 2715–2729.

Heinz, S., Benner, C., Spann, N., Bertolino, E., Lin, Y.C., Laslo, P., Cheng, J.X., Murre, C., Singh, H., and Glass, C.K. (2010). Simple combinations of lineage-determining transcription factors prime cis-regulatory elements required for macrophage and B cell identities. Mol Cell 38, 576–589.

Hnisz, D., Shrinivas, K., Young, R.A., Chakraborty, A.K., and Sharp, P.A. (2017). A Phase Separation Model for Transcriptional Control. Cell 169, 13–23.

Huang, Y., Sitwala, K., Bronstein, J., Sanders, D., Dandekar, M., Collins, C., Robertson, G., MacDonald, J., Cezard, T., Bilenky, M., et al. (2012). Identification and characterization of Hoxa9 binding sites in hematopoietic cells. Blood 119, 388–398.

Hyman-Walsh, C., Bjerke, G.A., and Wotton, D. (2010). An autoinhibitory effect of the homothorax domain of Meis2. FEBS J 277, 2584–2597.

Jacobs, J., Atkins, M., Davie, K., Imrichova, H., Romanelli, L., Christiaens, V., Hulselmans, G., Potier, D., Wouters, J., Taskiran, II, et al. (2018). The transcription factor Grainy head primes epithelial enhancers for spatiotemporal activation by displacing nucleosomes. Nat Genet 50, 1011–1020.

Jolma, A., Yin, Y., Nitta, K.R., Dave, K., Popov, A., Taipale, M., Enge, M., Kivioja, T., Morgunova, E., and Taipale, J. (2015). DNA-dependent formation of transcription factor pairs alters their binding specificity. Nature 527, 384–388.

Krumlauf, R. (1994). Hox genes in vertebrate development. Cell 78, 191–201.

Kutejova, E., Engist, B., Self, M., Oliver, G., Kirilenko, P., and Bobola, N. (2008). Six2 functions redundantly immediately downstream of Hoxa2. Development 135, 1463–1470.

Lambert, B., Vandeputte, J., Remacle, S., Bergiers, I., Simonis, N., Twizere, J.C., Vidal, M., and Rezsohazy, R. (2012). Protein interactions of the transcription factor Hoxa1. Bmc Dev Biol 12.

Langmead, B., and Salzberg, S.L. (2012). Fast gapped-read alignment with Bowtie 2. Nature methods 9, 357–359.

Lawrence, M., Huber, W., Pages, H., Aboyoun, P., Carlson, M., Gentleman, R., Morgan, M.T., and Carey, V.J. (2013). Software for computing and annotating genomic ranges. PLoS Comput Biol 9, e1003118.

Li, H., Handsaker, B., Wysoker, A., Fennell, T., Ruan, J., Homer, N., Marth, G., Abecasis, G., Durbin, R., and Genome Project Data Processing, S. (2009). The Sequence Alignment/Map format and SAMtools. Bioinformatics 25, 2078–2079.

Losa, M., Latorre, V., Andrabi, M., Ladam, F., Sagerstrom, C., Novoa, A., Zarrineh, P., Bridoux, L., Hanley, N.A., Mallo, M., et al. (2017). A tissue-specific, Gata6-driven transcriptional program instructs remodeling of the mature arterial tree. Elife 6.

Manley, N.R., and Capecchi, M.R. (1995). The role of Hoxa-3 in mouse thymus and thyroid development. Development 121, 1989–2003.

Manley, N.R., and Capecchi, M.R. (1997). Hox group 3 paralogous genes act synergistically in the formation of somitic and neural crest-derived structures. Dev Biol 192, 274–288.

McLean, C.Y., Bristor, D., Hiller, M., Clarke, S.L., Schaar, B.T., Lowe, C.B., Wenger, A.M., and Bejerano, G. (2010). GREAT improves functional interpretation of cis-regulatory regions. Nat Biotechnol 28, 495–501.

Merabet, S., and Mann, R.S. (2016). To Be Specific or Not: The Critical Relationship Between Hox And TALE Proteins. Trends Genet 32, 334–347.

Mirny, L.A. (2010). Nucleosome-mediated cooperativity between transcription factors. Proc Natl Acad Sci U S A 107, 22534–22539.

Moyle-Heyrman, G., Tims, H.S., and Widom, J. (2011). Structural constraints in collaborative competition of transcription factors against the nucleosome. Journal of molecular biology 412, 634–646.

Noyes, M.B., Christensen, R.G., Wakabayashi, A., Stormo, G.D., Brodsky, M.H., and Wolfe, S.A. (2008). Analysis of homeodomain specificities allows the family-wide prediction of preferred recognition sites. Cell 133, 1277–1289.

Pagès H, A.P., Gentleman R, DebRoy S (2019). Biostrings: Efficient manipulation of biological strings.

Pearson, J.C., Lemons, D., and McGinnis, W. (2005). Modulating Hox gene functions during animal body patterning. Nat Rev Genet 6, 893–904.

Porcelli, D., Fischer, B., Russell, S., and White, R. (2019). Chromatin accessibility plays a key role in selective targeting of Hox proteins. Genome Biol 20, 115.

Phuycharoen, M., Zarrineh, P., Bridoux, L., Amin, S., Losa, M., Chen, K., Bobola, N., and Rattray, M. (2019). Uncovering tissue-specific binding features from differential deep learning.

Reiter, F., Wienerroither, S., and Stark, A. (2017). Combinatorial function of transcription factors and cofactors. Curr Opin Genet Dev 43, 73–81.

Rezsohazy, R., Saurin, A.J., Maurel-Zaffran, C., and Graba, Y. (2015). Cellular and molecular insights into Hox protein action. Development 142, 1212–1227.

Rijli, F.M., Mark, M., Lakkaraju, S., Dierich, A., Dolle, P., and Chambon, P. (1993). A homeotic transformation is generated in the rostral branchial region of the head by disruption of Hoxa-2, which acts as a selector gene. Cell 75, 1333–1349.

Robinson, M.D., McCarthy, D.J., and Smyth, G.K. (2010). edgeR: a Bioconductor package for differential expression analysis of digital gene expression data. Bioinformatics 26, 139–140.

Ross-Innes, C.S., Stark, R., Teschendorff, A.E., Holmes, K.A., Ali, H.R., Dunning, M.J., Brown, G.D., Gojis, O., Ellis, I.O., Green, A.R., et al. (2012). Differential oestrogen receptor binding is associated with clinical outcome in breast cancer. Nature 481, 389–393.

Rual, J.F., Venkatesan, K., Hao, T., Hirozane-Kishikawa, T., Dricot, A., Li, N., Berriz, G.F., Gibbons, F.D., Dreze, M., Ayivi-Guedehoussou, N., et al. (2005). Towards a proteome-scale map of the human protein-protein interaction network. Nature 437, 1173–1178.

Selleri, L., Zappavigna, V., and Ferretti, E. (2019). ‘Building a perfect body’: control of vertebrate organogenesis by PBX-dependent regulatory networks. Genes Dev 33, 258–275.

Slattery, M., Riley, T., Liu, P., Abe, N., Gomez-Alcala, P., Dror, I., Zhou, T., Rohs, R., Honig, B., Bussemaker, H.J., et al. (2011). Cofactor binding evokes latent differences in DNA binding specificity between Hox proteins. Cell 147, 1270–1282.

Spitz, F., and Furlong, E.E. (2012). Transcription factors: from enhancer binding to developmental control. Nat Rev Genet 13, 613–626.

Staby, L., O’Shea, C., Willemoes, M., Theisen, F., Kragelund, B.B., and Skriver, K. (2017). Eukaryotic transcription factors: paradigms of protein intrinsic disorder. Biochem J 474, 2509–2532.

Tsai, A., Muthusamy, A.K., Alves, M.R., Lavis, L.D., Singer, R.H., Stern, D.L., and Crocker, J. (2017). Nuclear microenvironments modulate transcription from low-affinity enhancers. Elife 6.

Wickham (2016). Elegant Graphics for Data Analysis (Springer).

Zhang, Y., Liu, T., Meyer, C.A., Eeckhoute, J., Johnson, D.S., Bernstein, B.E., Nusbaum, C., Myers, R.M., Brown, M., Li, W., et al. (2008). Model-based analysis of ChIP-Seq (MACS). Genome Biol 9, R137.

