## Supplemental Figures with legends for "HOX paralogs selectively convert binding of ubiquitous transcription factors into tissue-specific patterns of enhancer activation"

### Figure 1- Figure Supplement 1

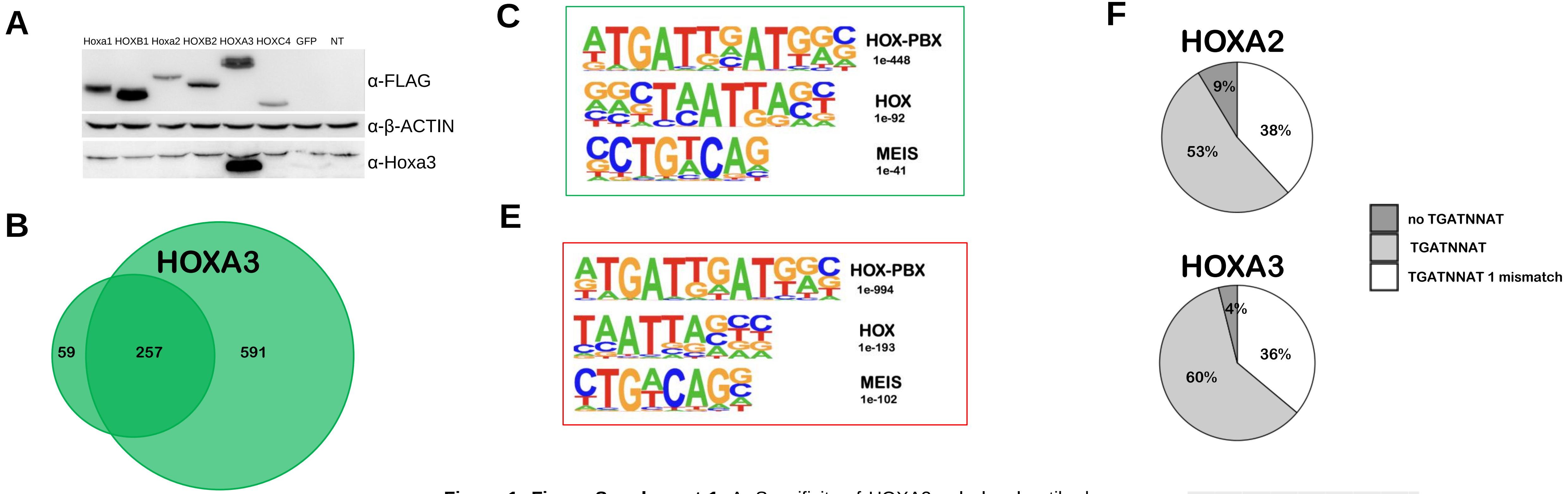


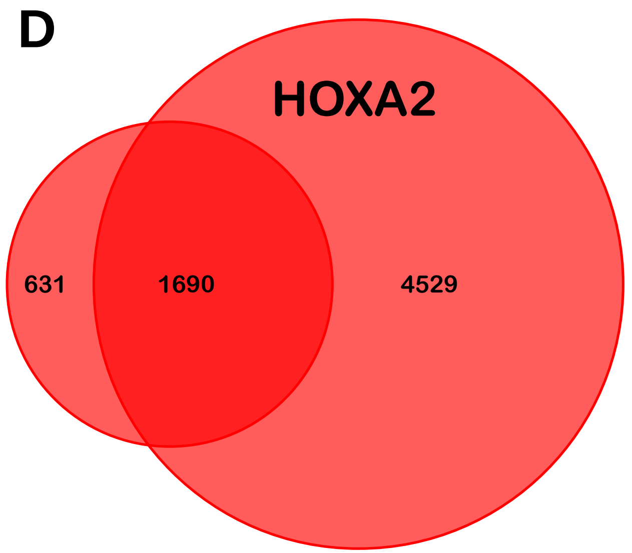


**Figure 1- Figure Supplement 1**. A. Specificity of HOXA3 polyclonal antibody. Western blot using FLAG and HOXA3 antibodies, as indicated. HOXA3 antibody recognizes HOXA3 and does not cross react with mouse HOXA1, HOXA2 and human HOXB1, HOXB2 and HOXC4. Plasmids containing *HOX*-FLAG cDNAs were transfected in HEK293 cells. B. Comparison of HOXA3 ChIP-seq replicates in E11.5 PBA (FE≥10). The replicate with larger genome coverage contains most (80%) of the second replicate's peaks and is used in all subsequent analyses. C. Top three most significant motifs identified in HOXA3 peaks using *de novo* motif discovery. D. Comparison of HOXA2 ChIP-seq replicates in E11.5 BA2. The smallest replicate was sequenced using the Illumina platform and is used for subsequent analysis; it largely (73%) overlaps HOXA2 ChIP-seq run on the Solid platform (Donaldson et al., 2012). E. Sequence logo of the top three most significant motifs identified in HOXA2 peaks. F. Frequency of HOX-PBX motifs in HOXA3 and HOXA2 peaks. G. Ratio of TGATNNAT motifs (perfect matches or 1nt mismatch allowed) in increasing numbers of HOXA3 (green) and HOXA2 (red) peaks, ordered by FE.

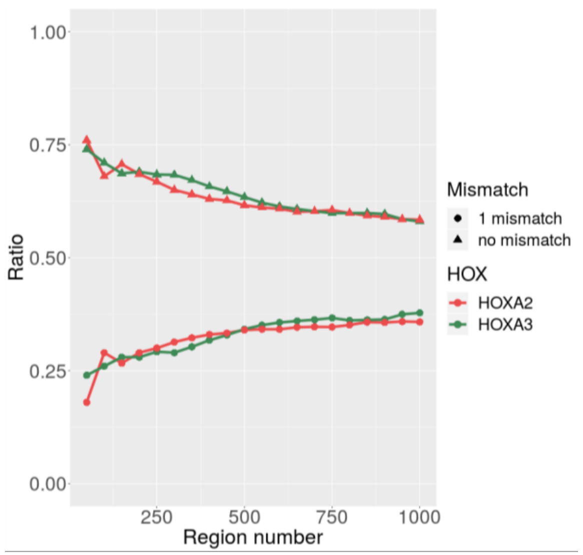

### Figure 2- Figure Supplement 1

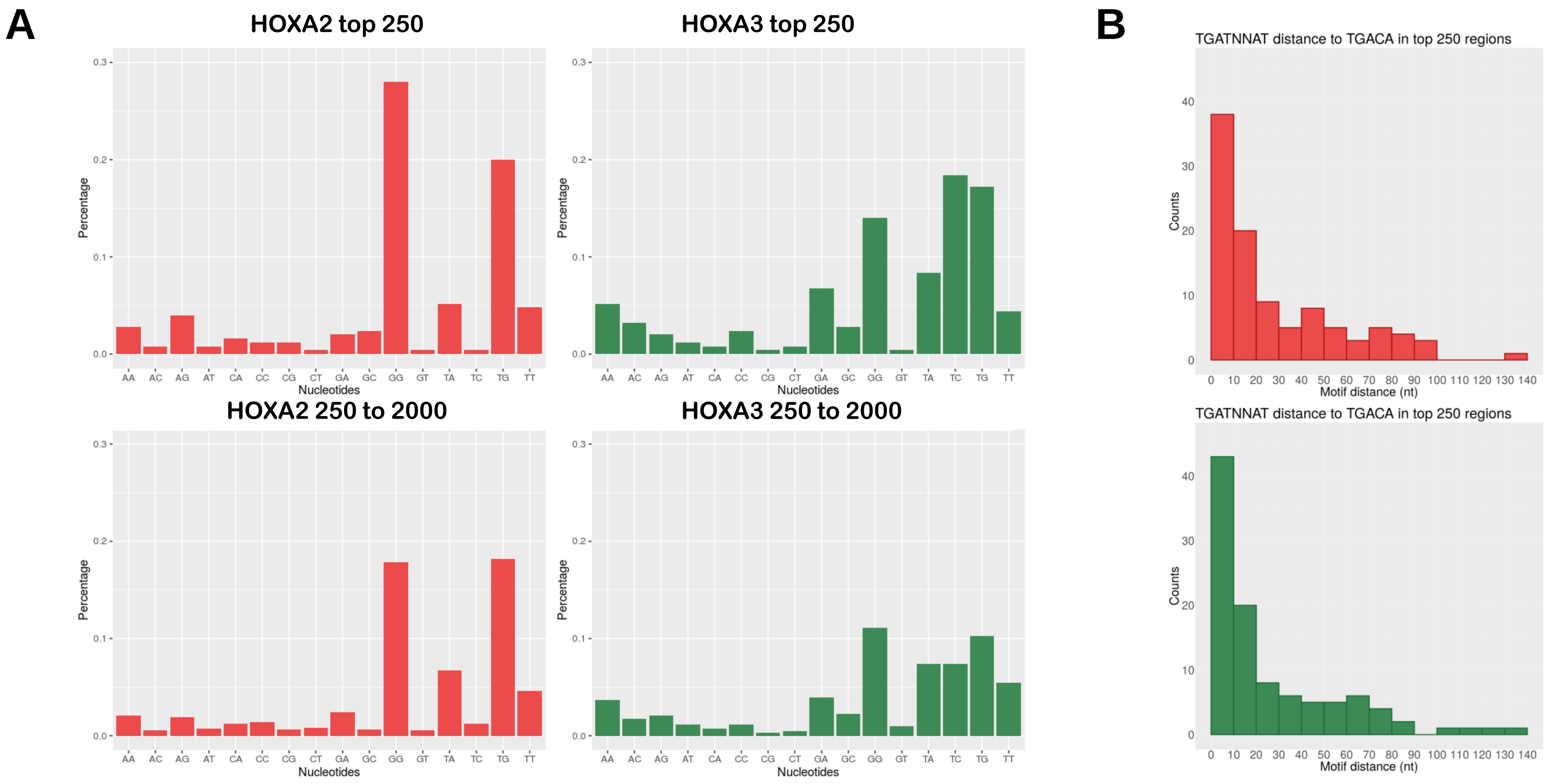


**Figure 2- Figure Supplement 1.** A. Percentage of TGATNNAT variants in top 250 or in top 250 to 2000 HOXA2 and HOXA3 peaks, as indicated. HOX-selective binding is more evident in the fraction of high-confidence peaks, suggesting that analysis of whole ChIP-seq experiments may mask effective HOX specificity. B. Distance between TGATNNAT (HOX-PBX) and TGACA (MEIS) in top 250 HOXA2 (red) and HOXA3 (green) peaks. Most TGACA occur at <20 nt from a HOX-PBX site.

### Figure 3- Figure Supplement 1

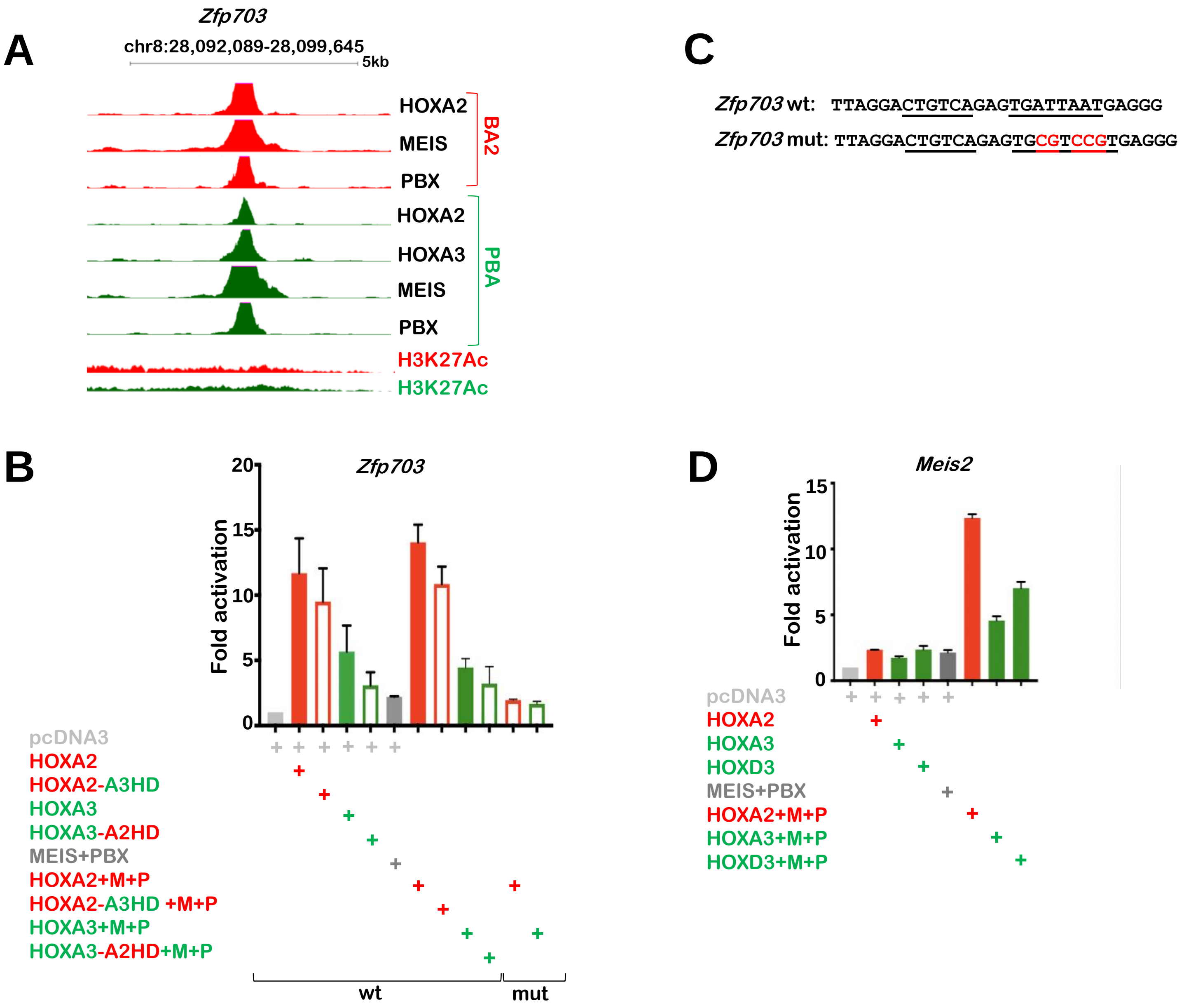


**Figure 3- Figure Supplement 1.** A. UCSC tracks of HOXA2, HOXA3, PBX, MEIS binding and H3K27 acetylation profiles in BA2 (red) and PBA (green) at the *Zfp703* locus. Strong HOX and TALE binding is observed in both tissues, with higher acetylation levels in BA2. B. Luciferase activity driven by *Zfp703* enhancer co-transfected with *Hoxa2* (red bar), *Hoxa3* (green bar), *Hoxa2-a3HD* (red empty bar), *Hoxa3-a2HD* (green empty bar), *Meis2* and *Pbx1a* (grey bar) expression vectors, alone and in combination, in NIH3T3 cells. The combination of *Hoxa2* with *Meis2* and *Pbx1a* results in the highest activation. Changing the HOX-PBX site (same nucleotide substitutions as in 3F) reduces HOX-TALE activation to the levels observed with TALE alone (empty bars). C. Sequence of *Zfp703* wild-type and mutant enhancer (only HOX-PBX and MEIS sites are shown). HOX-PBX and MEIS motifs are underlined. Nucleotide substitution in the HOX-PBX site are shown in red. D. Luciferase activity driven by *Meis2* enhancer co-transfected with *Hoxa2* (red bar), *Hoxa3* (green bar), *Hoxd3* (green bar), *Meis2* and *Pbx1a*, alone and in combination. When co-expressed with MEIS and PBX, HOXA2 shows higher activation capacity than HOX paralog 3. Values shown in BC represent fold activation over basal enhancer activity and are presented as the average of at least two independent experiments, each performed in triplicate. Error bars represent the SEM.

### Figure 5- Figure Supplement 1

A

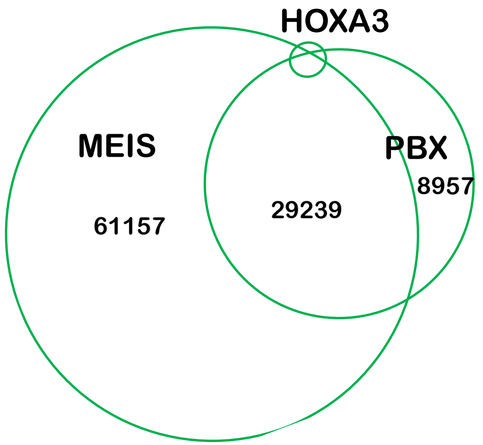


**Figure 5- Figure Supplement 1.** A. Whole Venn diagram of HOXA3, MEIS and PBX binding overlap, cropped in Fig. 5A. Most MEIS and PBX peaks do not overlap with any HOXA3 peaks. B-D. Kernel density plots of MEIS peaks relative to FE. MEIS binding is sorted into peaks not overlapping HOX (lighter colour) and peaks overlapping HOX as indicated (darker colour). HOX and MEIS ChIP-seq in mouse embryonic stem cells (HOXA1), BA2 (HOXA2) and hematopoietic stem cells (HOXA9) were used in this analysis.

B C D

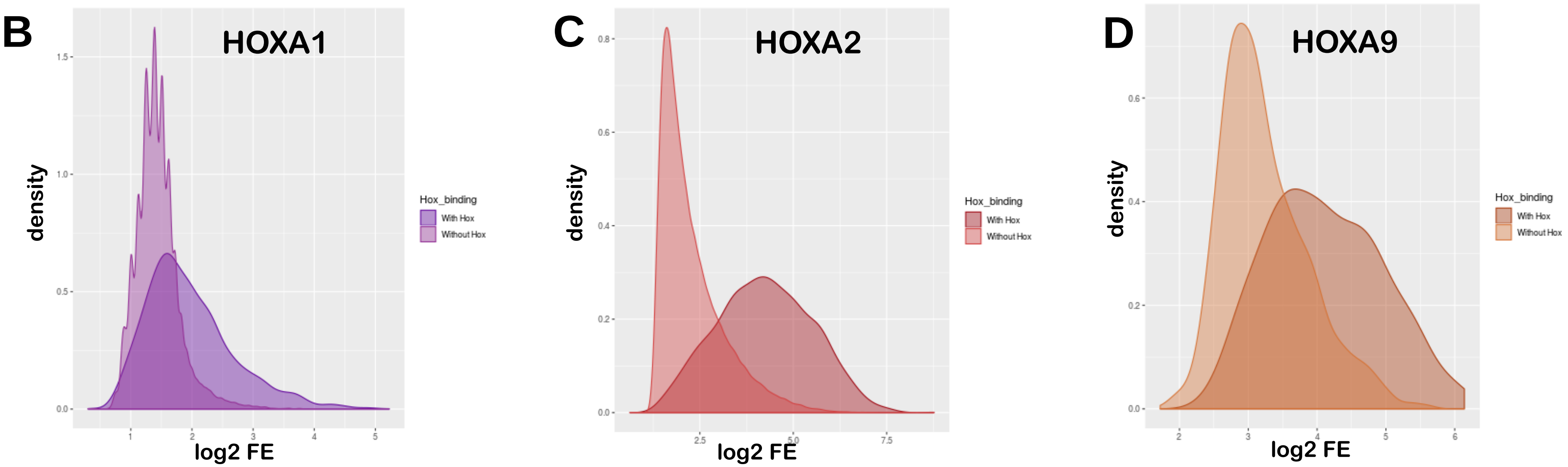

### Figure 6 - Figure Supplement 1

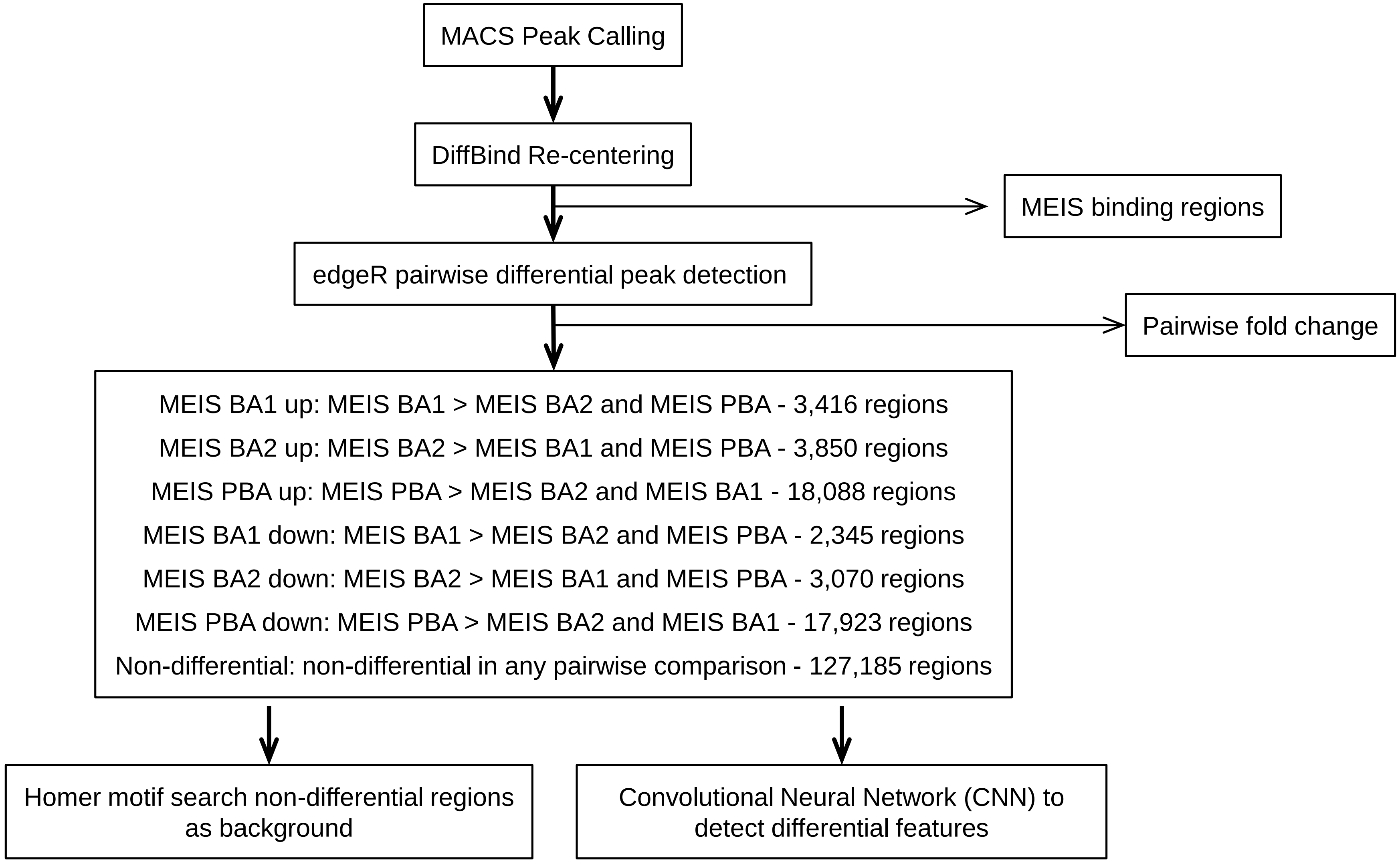


**Figure 6- Figure Supplement 1**. Identification of MEIS differential binding. MACS peaks from MEIS ChIP-seq experiments in BA1, BA2 and PBA with replicates were recentered using DiffBind (see methods), and pairwise fold changes and differential binding regions were computed by edgeR as described (Phuycharoen et. al. 2019). Pairwise differential regions were combined into MEIS BA1 up, BA1 down, BA2 up, BA2 down, PBA up, and PBA down regions. Differential sequence motifs were detected as described (Phuycharoen et. al. 2019). Differential H3K27ac regions and fold changes were computed in the same manner.

### Figure 6 - Figure Supplement 2

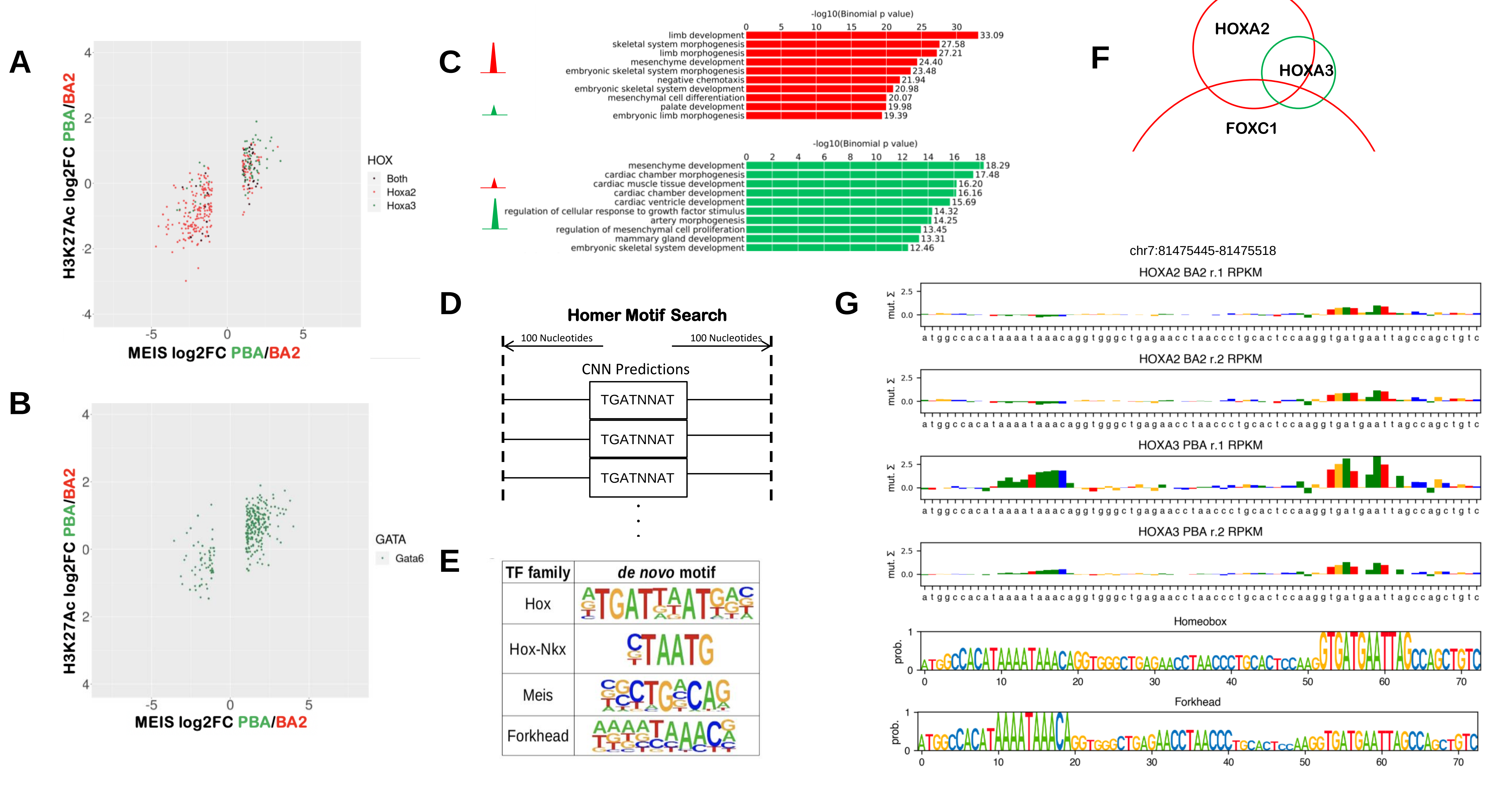

**Figure 6 - Figure Supplement 2**. A.B. Fold changes in MEIS binding and H3K27ac in PBA versus BA2 in HOX and GATA peaks. A. HOXA3 binding in PBA (green) and HOXA2 binding in BA2 (red) generally increases MEIS binding and H3K27ac in the corresponding tissues. B. Similarly, GATA6 binding in PBA increases MEIS and H3K27ac in PBA. C. GREAT analysis of MEIS peaks higher in BA2 (relative to PBA, red) and PBA (relative to BA2, green). Fold change ≥5 was used as a cutoff. Differential MEIS peaks are associated with genes involved in different biological processes (top 10 categories are shown). D.E. HOMER motif search in the vicinity of HOX/PBX sequence features identified in CNN MEIS higher PBA (+/-100 nt around TGATNNAT, as shown in D) identifies enrichment of Forkhead motifs. Other enriched flanking motifs include MEIS and HD motifs. F. Overlap of FOXC1 binding in BA2 (Amin et al., 2015), HOXA3 binding in PBA and HOXA2 binding in BA2 (red and green correspond to TF occupancy in BA2 and PBA, respectively). Only peaks with FE≥10 are considered. G. Example of HOX RPKM features, identified by CNN in a HOX-Forkhead bound region. The RPKM feature tracks show combined RPKM change caused by mutating each nucleotide to its alternatives. Features of two experimental replicates for HOXA2 in BA2 and two replicates of HOXA3 in PBA are predicted by a jointly trained multitask model, trained by transfer learning from a MEIS RPKM model. Below, normalised cross-correlation of region sequence with HOX and Forkhead position weight matrix (PWM) obtained by k-mer counting is shown. Nucleotides in HOX and Forkhead motifs clearly show increased sensitivity to perturbation, compared to their surrounding sequences.
